# High recallability of memory B cells requires ZFP318-dependent transcriptional regulation of mitochondrial function

**DOI:** 10.1101/2023.06.11.542551

**Authors:** Yifeng Wang, Wen Shao, Xin Liu, Qingtai Liang, Jiaqi Lei, Wenjuan Shi, Miao Mei, Ying Li, Xu Tan, Guocan Yu, Li Yu, Linqi Zhang, Hai Qi

## Abstract

Expression of the transcriptional regulator ZFP318 is induced in germinal center (GC)- exiting memory B cell precursors and memory B cells (MBCs). Using a conditional ZFP318 fluorescence reporter that also enables ablation of ZFP318-expressing cells, we found that ZFP318-expressing MBCs were highly enriched with GC-derived cells. Although ZFP318-expressing MBCs constituted only a minority of the antigen-specific MBC compartment, their ablation severely impaired recall responses. Deletion of Zfp318 did not alter the magnitude of primary responses but markedly reduced MBC participation in recall. CD40 ligation promoted Zfp318 expression, whereas B-cell receptor (BCR) signaling was inhibitory. Enforced ZFP318 expression enhanced recall performance of MBCs that otherwise responded poorly. ZFP318-deficient MBCs expressed less mitochondrial genes, had structurally compromised mitochondria, and were susceptible to reactivation-induced cell death. The abundance of ZFP318-expressing MBCs, instead of the number of antigen-specific MBCs, correlated with the potency of prime-boost vaccination. Therefore, ZFP318 controls the MBC recallability and represents a quality checkpoint of humoral immune memory.

## Introduction

Memory B cells (MBCs) play a key role in vaccine-induced protective immunity against infectious diseases. In a standard prime-boost vaccine regimen, primary immunization leads to the development of MBCs that, upon boost, are re-activated to produce long-lived plasma cells (LLPCs) to sustain serum antibody titers for an extended period of time^1–3^. Some re-activated MBCs participate in secondary germinal center (GC) reaction^4–7^, and by undergoing secondary diversification and selection they may lead to emergence of antibodies that recognize pathogen variants and antibodies that can broadly neutralize mutating viruses^8,9^. Therefore, successful prime-boost vaccines depend on high-quality functional MBCs developed during the prime. Extensive efforts have been devoted to understanding how MBCs develop and how subsets of MBCs may preferentially produce secondary GCs or PCs in the recall response^1,10,11^. However, adoptive transfer of a large number of input MBCs is seemingly not sufficient for generating a robust output of donor-derived GCs or PCs ^12,13^, in sharp contrast to what is commonly observed with memory T cells. A provocative possibility is that some antigen-specific MBCs may not be intrinsically recallable, that is, capable of being reactivated by the same antigen to participate in a productive response.

We and others have previously reported identification of GC memory precursors^13–15^. GC light-zone B cells that have reached the G0 phase, as measured using a cell cycle-reporting system, can functionally reconstitute a secondary response and express a transcriptome similar to that of MBCs^13^. Ephrin-B1^+^S1PR2^lo^ or the CCR6^+^ fraction of GC light-zone cells are enriched with cells expressing MBC-resembling transcriptomes^14,15^. By different approaches, these studies report very similar transcriptomic features of GC memory precursors. Presumably, expression of genes that are uniquely important for MBC but not for GC functions would be increased already at the precursor stage. In looking for such genes, we noticed that GC memory precursors increased expression of *Zfp318*, a gene coding for a transcription regulator that is essential for spermatogenesis^16^ and IgD expression in naïve B cells^17,18^. ZFP318 promotes IgD expression by repressing recognition of the transcription termination site upstream of the coding sequence for the IgD heavy chain^18^. However, by definition IgD is not expressed by GC memory precursors and MBCs. Therefore, we asked whether ZFP318 might play a specific role in regulating MBC development and/or function, distinct from its role in the regulation of IgD expression.

## Results

### Characterization of the ZSDAT reporter allele for the *Zfp318* gene

We first sought to develop a loss-of-function allele that can also report transcription activity of the *Zfp318* locus in a cell type-specific manner. Because MBCs are heterogenous in phenotype and function, we wanted also to build into the loss-of-function reporter allele a cell-ablation element for inducible deletion of ZFP318-expressing cells. Therefore, immediately downstream of the transcription start site of the *Zfp318* gene, we inserted a construct containing a STOP cassette flanked by two loxP sites, which is followed sequentially by the coding sequence for diphtheria toxin receptor (DTR), the coding sequence for the self-cleaving 2A peptide of *Thosea asigna* virus, and the coding sequence for tdTomato (Fig.1A). This compound mutant allele cannot express *Zfp318* gene, but it permits expression of both DTR and tdTomato under the endogenous *Zfp318* locus control after excision of the STOP cassette by a Cre driver. We denote this modified allele ZSDAT (*<u>Z</u>fp318*-loxP-<u>S</u>TOP-loxP-<u>D</u>TR-2<u>A</u>-td<u>T</u>omato), and *Zfp318*^ZSDAT/+^ represents a heterozygous genotype that carries a wildtype *Zfp318* allele able to normally express the gene and a mutant ZSDAT allele that cannot express *Zfp318* but reports its transcription activity.

**Figure 1.**
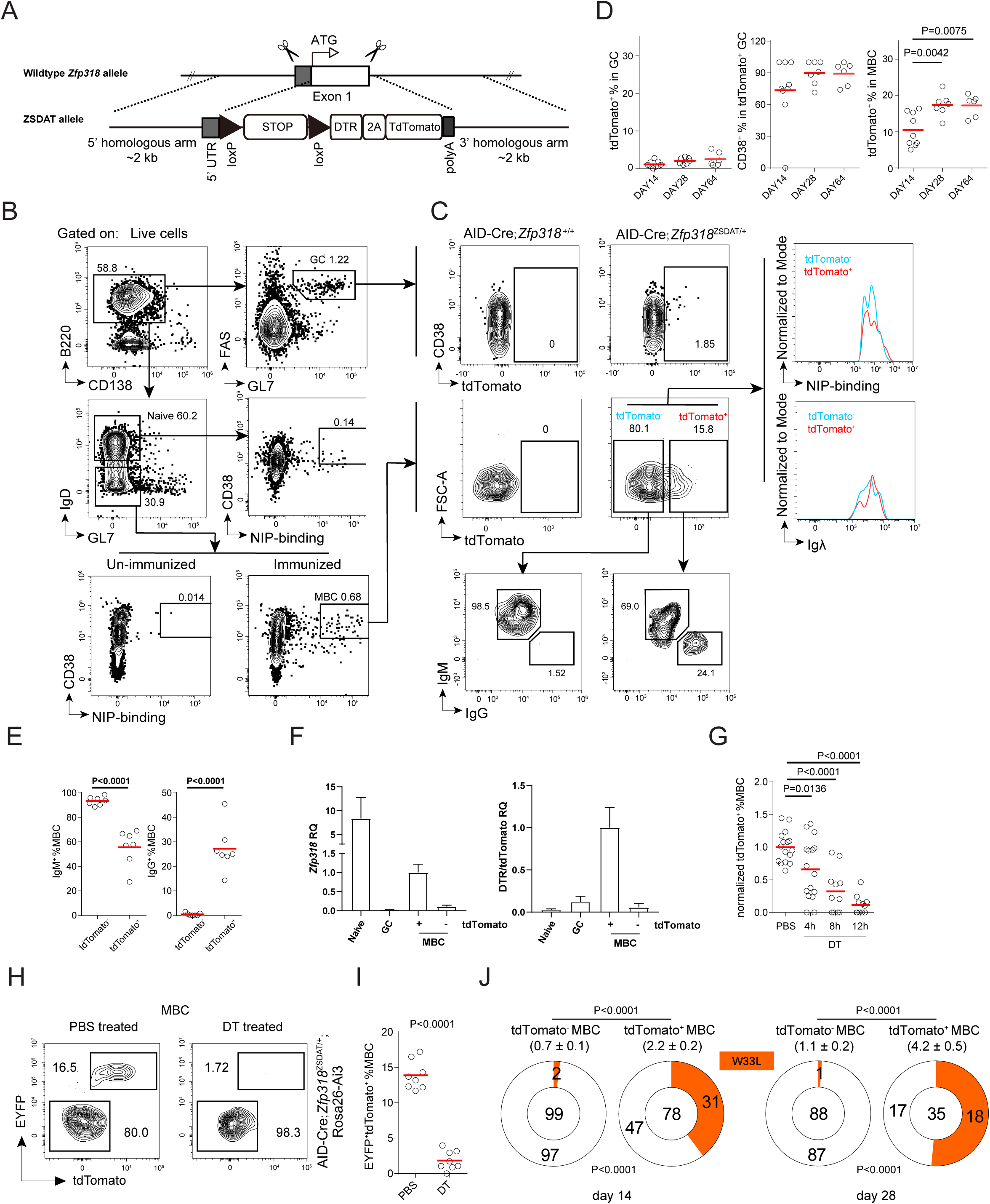
MBC heterogeneity revealed by the ZSDAT reporter. **(A)** The diagram of ZSDAT reporter allele. STOP, transcription stop cassette; black triangles, flox sites. **(B)** The gating strategy for sorting GC (top), naïve B cells (middle) and NIP-binding MBCs (bottom). The lower boundary for the NIP-binding gate of MBCs is set according to unimmunized control, and the lower boundary for the CD38 gate of MBCs is based on the upper boundary of GCs. **(C)** Representative tdTomato fluorescence profiles of GCs and NIP-binding MBCs from AID-Cre or AID-Cre;*Zfp318*^ZSDAT/+^ mice 14 days after NP-KLH immunization. MBCs in AID-Cre;*Zfp318*^ZSDAT/+^ mice were further separated into tdTomato^+^ and tdTomato^-^ cells, with their respective isotype composition (bottom), NIP binding and Igλ expression profiles (right) shown as indicated. **(D)** Frequencies of tdTomato^+^ cells in GCs, and MBCs observed 14, 28 or 64 days after NP-KLH immunization. Data pooled from 2 independent experiments. *P* values by unpaired student t-test. **(E)** Frequencies of IgM^+^ (left) or IgG^+^ (right) cells observed in tdTomato^+^ or tdTomato^-^ MBCs 14 days post NP-KLH immunization. **(F)** ZFP318 (top panel) and corresponding DTR/tdtomato mRNA levels (bottom panel) by quantitative RT-PCR in tdTomato^+^ or tdTomato^-^ cells of indicated types (GFP-expressing AID-Cre;*Zfp318*^ZSDAT/+^ B1-8^hi^ cells in B6 mice 7 days after NP-OVA immunization). **(G)** Normalized abundance of the tdTomato^+^ fraction in MBCs from NP-KLH-immunized AID-Cre;*Zfp318*^ZSDAT/+^ mice, at indicated time points after DT treatment. Data were pooled from 4 independent experiments. In each, the mean value of the PBS group is set as unit 1. (**H-I**) Representative FACS profiles of EYFP (Rosa26-Ai3) and tdTomato (ZSDAT) fluorescence (**H**) and summary data (**I**) of MBCs AID-Cre;*Zfp318*^ZSDAT/+^;Rosa26-Ai3 mice treated with PBS (left) or DT (right) for 24 hours that are 60 days after NP-KLH immunization. **(J)** Overall mutations (mean±s.e.m.) per V_H_186.2 heavy chain and frequencies of affinity-enhancing W33L mutations in tdTomato^+^ or tdTomato^-^ NIP-binding MBCs sorted from AID-Cre;*Zfp318*^ZSDAT/+^ mice 14 or 28 days after immunization. Numbers of analyzed clones are indicated at the center of the pie chart. Data are pooled from 2 independent experiments, involving >10 mice per group. *P* values by *t* tests (top, overall mutations) and by Fisher’s exact test (bottom, W33L mutations).

The AID-Cre line can track post-activation B cells that previously expressed AID, mainly those that are derived from GCs ^19^. Naïve AID-Cre;*Zfp318*^ZSDAT/+^ B cells do not express AID and thus would have no tdTomato fluorescence, whereas GC-derived MBCs should become tdTomato^+^ if their *Zfp318* locus is transcriptionally active. On the other hand, tdTomato^-^ MBCs would contain GC-derived MBCs that do not express *Zfp318* and other MBCs that have never expressed AID, regardless of whether their *Zfp318* locus is active or not.

To characterize the system, we immunized AID-Cre;*Zfp318*^ZSDAT/+^ mice with NP-KLH. The replacement of one wildtype *Zfp318* allele with the ZSDAT allele did not alter the primary response to NP-KLH immunization, as evidenced by the similar GC, SPPC, BMPC and NIP-specific MBC abundance in AID-Cre;*Zfp318*^ZSDAT/+^ and control AID-Cre;*Zfp318*^+/+^ mice and by similar titers of NP-specific antibodies of comparable affinities in these two groups of mice (Fig. S1). Naïve B cells from AID-Cre;*Zfp318*^ZSDAT/+^ mice expressed no tdTomato (not shown). As shown in Fig. 1B-D, the bulk of Fas^+^GL7^+^ GC B cells contained only a very small fraction of tdTomato^+^ cells, most of which were CD38^+^, consistent with the increased ZFP318 expression in GC memory precursors that began to express CD38. NIP-binding MBCs contained a larger fraction of tdTomato^+^ cells, ranging from ∼10% at day 14 to ∼20% at day 28 and later (Fig. 1D). Nonetheless, tdTomato^+^ cells constituted a minority in all NIP-binding MBCs, and they were ∼25% IgG^+^ whereas tdTomato-MBCs were mostly IgM^+^ (Fig. 1C, bottom; quantitation in Fig. 1E).

To further validate the system, we took an adoptive transfer approach in order to acquire sufficient numbers of MBCs for quantitative analysis of *Zfp318* expression *per se* by RT-PCR. To this end, CD45.2 AID-Cre;*Zfp318*^ZSDAT/+^ B1-8^hi^ BCR-transgenic cells were transferred into CD45.1 recipients. Following immunization of recipient mice, we sort-purified CD45.2 naïve, GC and tdTomato^+^ or tdTomato^-^ NP-binding MBC cells derived from donor B1-8^hi^ cells. As shown in Fig. 1F, naïve B cells expressed a high level of ZFP318 but no DTR, while bulk GC cells (containing the CD38^+^ fraction) expressed little ZFP318 or DTR; tdTomato^+^ MBCs expressed both ZFP318 and DTR, while tdTomato^-^ MBCs expressed little ZFP318 or DTR. Of note, tdTomato^-^ MBCs should theoretically contain cells that can currently express ZFP318 but have not expressed AID and therefore do not express tdTomato. However, little ZFP318 was detected in those tdTomato^-^ MBCs (Fig. 1E), indicating that such cells are very rare, if present at all. Overall, these data show that the tdTomato fluorescence in MBCs of the AID-Cre;*Zfp318*^ZSDAT/+^ reporter strain faithfully reports ZFP318 and DTR expression as designed. Consistent with DTR expression being restricted to tdTomato^+^ cells, treatment *in vivo* with diphtheria toxin (DT) led to deletion of the vast majority of tdTomato^+^ MBCs in 12 h (Fig. 1G).

We further bred AID-Cre;*Zfp318*^ZSDAT/+^ with the Rosa26-Ai3 reporter line to introduce an independent marker for tracking occurrence of AID expression. The majority of EYFP^+^ NIP-specific MBCs, those that had previously expressed AID, were tdTomato^+^ (Fig. 1H). EYFP^-^ MBCs, those that had not expressed AID, were tdTomato^-^, as expected. DT treatment could readily delete tdTomato^+^ cells (Fig. 1H-I). The NIP-binding and Igλ expression profiles of tdTomato^+^ and tdTomato^-^ MBCs were similar (Fig. 1C). The overall Ig somatic mutation load in tdTomato^+^ MBCs increased from 2.2±0.2 on day 14 to 4.2±0.5 on day 28, whereas the mutation load in tdTomato^-^ MBCs were low and minimally changed over time (0.7±0.1 to 1.1±0.2; Fig. 1J). At both time points, tdTomato^+^ MBCs harbored a higher frequency of W33L mutation that enhances the V_H_186.2 antibody affinity toward the NP hapten.

Taken together, these data demonstrate that the ZSDAT reporter allele performs as designed. They also indicate that tdTomato^+^ MBCs, expressing ZFP318, are enriched within GC-derived MBCs, whereas tdTomato^-^ MBCs are comparatively enriched within GC-independent MBCs.

### ZFP318-expressing MBCs are main contributors to antigen recall responses

To investigate the biological significance of heterogeneous ZFP318 expression among MBCs, we first took advantage of the ability to ablate ZFP318-expressing cells with diphtheria toxin (DT) afforded by the ZSDAT allele. Two months after primary immunization with NP-KLH, AID-Cre;*Zfp318*^ZSDAT/+^ mice were treated with either DT or PBS and then re-challenged (Fig. 2A). Six days later, a marked reduction in the numbers of NP-binding GCs, SPPCs, and BMPCs was seen in the DT-treated as compared to the PBS group (Fig. 2B-C). Importantly, whereas serum NP-specific titers rapidly increased in the PBS group (by ∼4 fold) in these 6 days, very little post-boost increase was seen in the DT group (Fig. 2D). Because DT at the dose applied had no impact on immune response in B6 mice (not shown), and ZFP318-expressing MBCs only accounted for ∼20% of the entire antigen-binding MBC compartment (Fig. 1D), these cells appear to have a disproportionately large contribution to the recall response.

**Figure 2.**
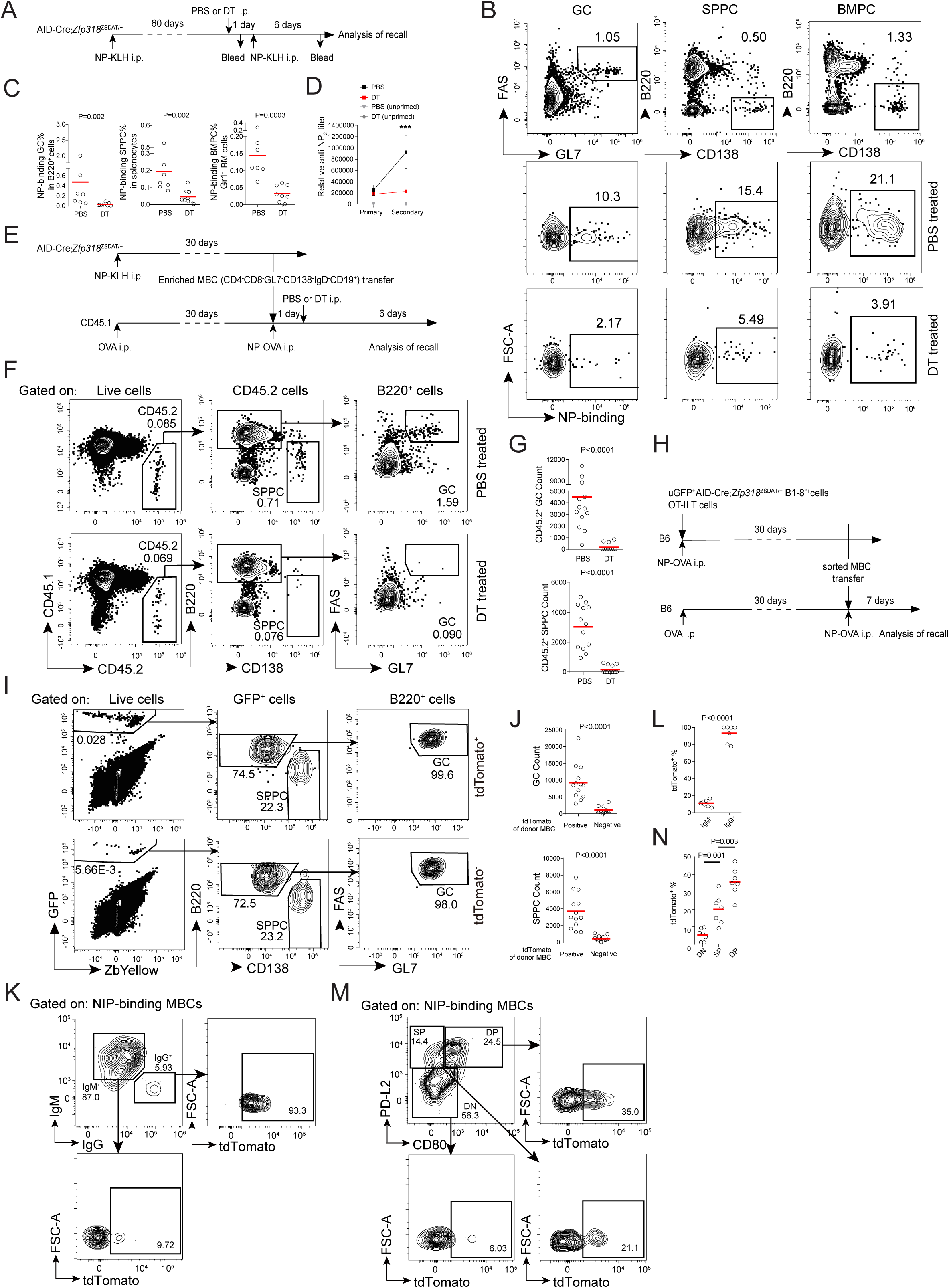
Recallable memory is carried in ZFP318-expressing MBCs. **(A-D)** Recall in AID-Cre;*Zfp318*^ZSDAT/+^ mice. **(A)** Experimental design. **(B-C)** Representative FACS profiles (**B**) and summary data (**C**) of GCs, splenic PCs, BMPCs in PBS- or DT-treated group 7 days after re-challenge, as done according to (**A**). Data were pooled from 2 independent experiments, with each symbol indicating one mouse and lines denoting means. *P* values by *t* tests. **(D)** Anti-NP_2_ IgG titers by ELISA. Unimmunized and challenged without prior immunization (unprimed) mice were used as controls. Data were pooled from 2 independent experiments. *P* values by t tests. **(E-G)** Recall in CD45.1 recipients of MBCs from AID-Cre;*Zfp318*^ZSDAT/+^ mice. (**E**) Experimental design. **(F-G)** Gating strategy to identify donor-derived cells (**F**) and summary data (**G**) of donor-derived GCs, splenic PCs, in PBS- or DT-treated group 7 days after re-challenge, as done according to (**E**). Data were pooled from 4 independent experiments, with each symbol indicating one mouse and lines denoting means. *P* values by *t* tests. **(H-J)** Recall in B6 recipients of B1-8^hi^ MBCs from uGFP-expressing AID-Cre;*Zfp318*^ZSDAT/+^ mice. **(H)** Experimental design. **(I-J)** Gating strategy to identify donor-derived cells (**I**) and summary data (**J**) of donor-derived GCs, splenic PCs, in tdTomato^+^ and tdTomato^-^ group 7 days after re-challenge, as done according to (**H**). Data were pooled from 4 independent experiments, with each symbol indicating one mouse and lines denoting means. *P* values by *t* tests. **(K-N)** ZFP318 expression in MBC subsets classified according to isotype or CD80 and PDL2 expression. **(K-L)** FACS profiles (**K**) and summary data (**L**) of tdTomato expression in IgG^+^ and IgM^+^ MBCs from mice of an AID-Cre;*Zfp318*^ZSDAT/+^ genotype 28 days post immunization. **(M-N)** FACS profiles (**M**) and summary data (**N**) of tdTomato expression in DP, SP and DN MBCs from mice of an AID-Cre;*Zfp318*^ZSDAT/+^ genotype 28 days post immunization. Data were pooled from 2 independent experiments, with each symbol indicating one mouse and lines denoting means. *P* values by *t* tests.

ZFP318 may positively regulate the recall performance of individual MBCs in a cell-intrinsic manner. However, because tdTomato^-^ MBCs enrich those derived from a GC-independent source and of lower affinities, it is possible that tdTomato^-^ MBCs were competitively inhibited by cells of higher affinities or by antibody feedback ^20^. Therefore, we measured recall performance of MBCs after adoptive transfer into naïve hosts. Splenocytes were obtained from AID-Cre;*Zfp318*^ZSDAT/+^ mice 30 days after NP-KLH immunization and subjected to magnetic depletion of CD4^+^ T cells, CD8^+^ T cells, IgD^+^ naïve B cells, GL7^+^ GC B cells, and CD138^+^ PCs. From such T-GC-PC-depleted splenocytes, CD19^+^ B cells were then positively selected with magnetic beads. In the resultant cell preparation, ∼40% cells were B220^+^ B cells of an IgD^-^GL7^-^CD138^-^ phenotype, containing essentially no naïve B cells, no GCs and no PCs but still contained NIP-binding MBCs (Fig. S2A). We transferred this cell preparation into CD45.1 recipient mice so that each recipient received an equal number of ∼2000 NIP-binding B220^+^CD38^+^IgD^-^ MBCs. Those recipient mice were primed with ovalbumin immunization 30 days earlier and re-immunized with NP-OVA on the day of MBC transfer. One day after MBC transfer, recipient mice were treated with PBS as control or DT to delete donor ZFP318-expressing MBCs (Fig. 2E). Donor MBCs gave rise to both secondary GCs and PCs upon recall in the control but very few in the DT-treated group (Fig. 2F-G), indicating that, among all NIP-binding MBCs, ZFP318-expressing MBCs are the predominant contributor to the recall response.

These observations with adoptive transfer support the possibility that MBCs are intrinsically different in their response to antigen re-challenge with varying efficiencies dependent on ZFP318. However, because tdTomato^+^ ZFP318-expressing MBCs carry BCR of higher affinities than tdTomato^-^ counterparts, ZFP318-expressing MBCs may respond better mainly because of higher affinities for antigen. Therefore, we compared MBCs that carry identical high-affinity NP-specific BCRs but differ in ZFP318 expression. To this end, we conducted adoptive transfer experiments using AID-Cre;*Zfp318*^ZSDAT/+^ B1-8^hi^ cells that also express human ubiquitin promoter-driven transgenic GFP (uGFP)^21^. Such B1-8^hi^ cells were co-transferred together with OT-II T cells into B6 mice, which were then immunized with NP-OVA (Fig. 2H). A month later, ∼60% of uGFP-expressing, donor-derived CD38^+^IgD^-^GL7^-^ NP-binding B1-8^hi^ MBCs were tdTomato^+^ (Fig. S2B). We sort-purified tdTomato^-^ and top 1/3 tdTomato^+^ NP-binding B1-8^hi^ MBCs (Fig. S2B) and transferred ∼2000 cells into each new B6 recipient that was primed with ovalbumin before (Fig. 2H). A week after NP-OVA immunization, tdTomato^+^ but not tdTomato^-^ B1-8^hi^ MBCs produced substantial amounts of secondary GCs and PCs (Fig. 2I-J). In this system, tdTomato^+^ and tdTomato^-^ B1-8^hi^ MBCs had essentially the same BCR and were mostly IgG^+^ (Fig. S2C) but differed in recall performance. These data suggest that tdTomato^+^ MBCs are superior in antigen recall because they are intrinsically different in a ZFP318-dependent manner.

### Relationship of ZFP318-expressing MBCs with regards to other phenotypically classified MBC subsets

It has been reported that recall generation of secondary GCs is favored by unswitched or PDL2^-^CD80^-^ MBCs, whereas recall generation of secondary PCs is favored by isotype-switched or PDL2^+^CD80^+^ MBCs ^22–24^. We found that IgM^+^ unswitched MBCs contained ∼10% tdTomato^+^ ZFP318-expressing cells, whereas ∼90% of IgG-switched MBCs were ZFP318-expressing (Fig. 2K-L). CD80^+^PDL2^+^, CD80^-^ PDL2^+^ and CD80^-^PDL2^-^ subsets contained ∼40%, 20% and <10% ZFP318-expressing cells (Fig. 2M-N). These data also imply that, on a per cell basis, IgG-switched MBCs are more functionally recallable than unswitched MBCs, and CD80^+^PDL2^+^ are more functional than CD80^-^PDL2^-^ MBCs.

### ZFP318 is required for the secondary but not primary antibody response

To test whether ZFP318 itself is required for optimal antigen recall performance of MBCs, we bred the ZSDAT knockout-reporter allele to homozygosity to create ZFP318-deficient mice. Consistent with the previous report of normal B cell development in the complete absence of ZFP318 (ref. ^18^), we observed no gross defects in B cell populations in AID-Cre;*Zfp318*^ZSDAT/ZSDAT^ knockout mice (data not shown). Moreover, as compared to AID-Cre;*Zfp318*^+/+^ control mice, they showed no defects in the primary response, producing comparable GCs, MBCs, PCs and NP-specific IgG titers after NP-KLH immunization (Fig. S3). Therefore, ZFP318 is not required for the primary B cell response and not required for MBC development. In sharp contrast, whereas a marked increase in NP-specific serum IgG titers was seen in control mice following NP-KLH re-challenge, as expected of a normal secondary response, ZFP318-deficient mice failed to do so (Fig. 3A-B). Therefore, the *Zfp318* gene function is required for the secondary but not for the primary antibody response, consistent with the possibility that ZFP318 controls MBC recall performance in a cell-intrinsic manner.

**Figure 3.**
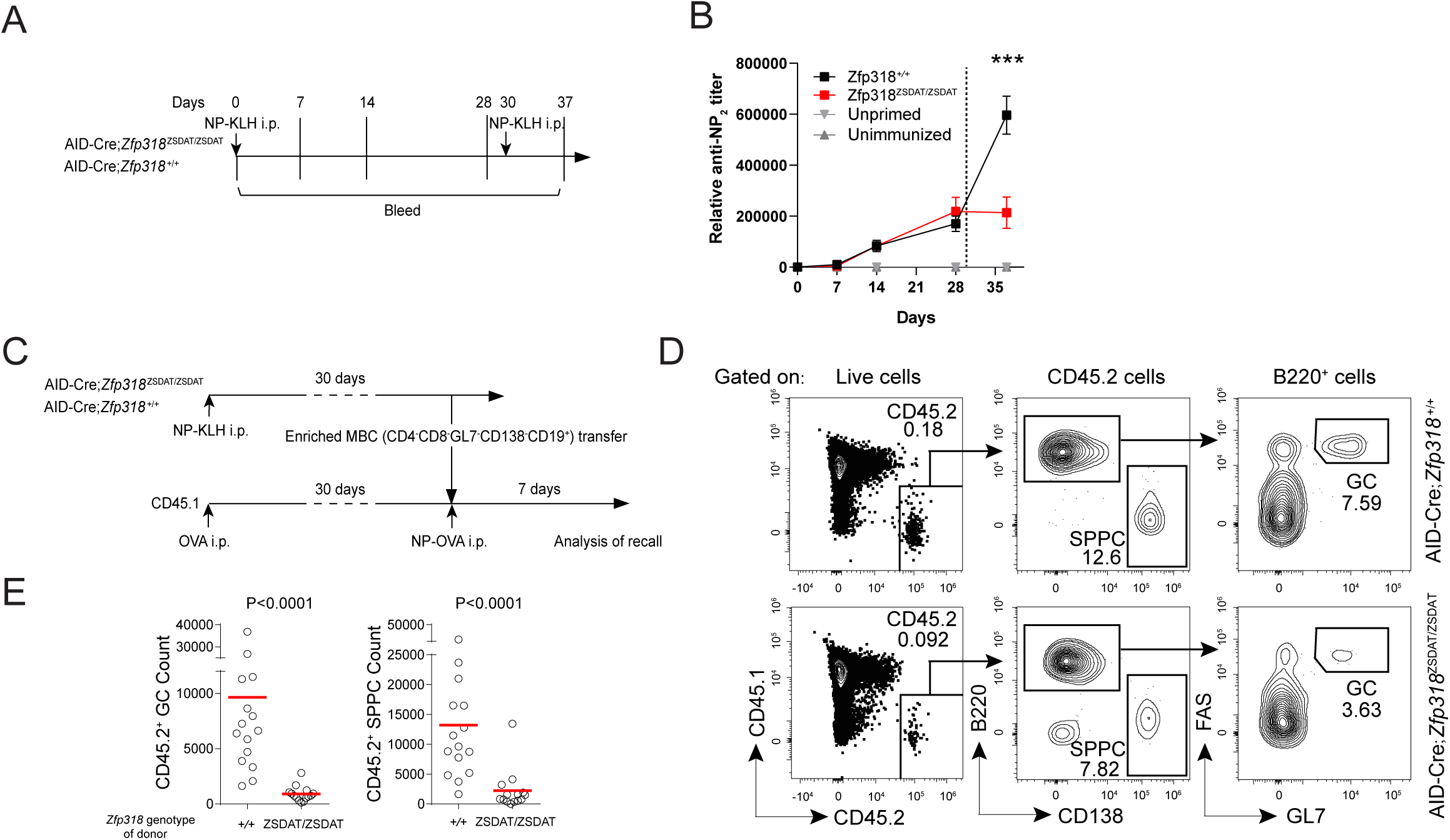
ZFP318 underlies MBC recallability. **(A-B)** Recall in AID-Cre;*Zfp318*^+/+^ and AID-Cre;*Zfp318*^ZSDAT/ZSDAT^ mice. (**A)** Experimental design. (**B)** Antibody titers of anti-NP_2_ IgG by ELISA. Dotted line indicates secondary challenge. Unimmunized and challenged without prior immunization (unprimed) mice were used as controls. Data were pooled from 2 independent experiments. *P* values by t tests. **(C-E)** Recall in CD45.1 recipients of MBCs from AID-Cre;*Zfp318*^+/+^ and AID-Cre;*Zfp318*^ZSDAT/ZSDAT^ mice. **(C)** Experimental design. **(D)** Gating strategies to identify GCs and SPPCs. **(E)** Summary data of cell abundance of indicated types. Data were pooled from 2 independent experiments, with each symbol indicating one mouse and lines denoting means. *P* values by Mann-Whitney tests.

To determine whether it is ZFP318 expressed in MBCs that is required for the MBC recall, we conducted MBC transfer and recall experiments. A month after NP-KLH immunization, NIP-binding MBCs were enriched from AID-Cre;*Zfp318*^ZSDAT/ZSDAT^ or control AID-Cre;*Zfp318*^+/+^ mice (Fig. 3C) as T/GC/PC-depleted CD19^+^ cells using essentially the same protocol as in Fig. 2E and Fig. S2A, except for that no IgD dumping was utilized because ZFP318-deficient B cells do not express IgD (ref. 17 and data not shown). The cell preparation was analyzed by cytometry to quantitate the abundance of NIP-binding MBCs, and ∼2000 cells of each genotype were transferred into CD45.1 OVA-primed recipient mice. Seven days after NP-OVA immunization of these CD45.1 recipients, whereas significant numbers of wildtype donor-derived secondary GCs and PCs were generated, very few ZFP318-deficient GCs or PCs were produced (Fig. 3D-E). The lack of IgD expression in naïve B cells could not explain the diminished recall by ZFP318-deficient MBCs, because MBCs derived from *Ighd*^-/-^ or wildtype naïve B cells mounted comparable recall responses (Fig. S3D-F). This finding echoes previous observations that an *Igd* deficiency has almost no impact on the secondary response^25^. Together, these results indicate that ZFP318 expressed by MBCs is required for these cells to mount a proper secondary response to antigen.

### Intrinsic requirement of ZFP318 for proper recall is independent of isotype or affinity variation

To rule out isotype or affinity variations between wildtype and ZFP318-deficient MBCs as a potential explanation for why ZFP318-deficient MBCs perform poorly in recall, we further compared recall performance of ZFP318-deficient (AID-Cre;*Zfp318*^ZSDAT/ZSDAT^) and ZFP318-heterzygous (AID-Cre;*Zfp318*^ZSDAT/+^) B1-8^hi^ MBCs, using the adoptive transfer protocol as outlined in Fig. S4A (essentially the same protocol as described in Fig. 2H and Fig. S2B). To this end, uGFP-expressing donor-derived NP-binding, tdTomato^+^ B1-8^hi^ MBCs were sort-purified 30 days after immunization, transferred at ∼2000 cells per mouse into a new set of OVA-primed B6 recipients and recalled by NP-OVA immunization. As shown in Fig. S4B, both ZFP318-deficient and -heterozygous B1-8^hi^ MBCs were predominantly IgG^+^. Inclusion of tdTomato^+^ as an additional sorting marker is to maximize the comparability of MBCs of the two genotypes: those ZFP318-deficient tdTomato^+^ MBCs were ZFP318-want-to-express cells with an active *Zfp318* locus that cannot express the *Zfp318* gene itself, while those ZFP318-heterozygous tdTomato^+^ MBCs do express. Seven days after recall, secondary GCs and PCs generated by ZFP318-deficient tdTomato^+^ MBCs were significantly diminished as compared to those generated by ZFP318-heterozygous tdTomato^+^ MBCs (Fig. S4C-D). These data rule out the possibility that affinity or isotype variation could account for the poor recall performance exhibited by MBCs that cannot express ZFP318, indicating ZFP318 itself as essential for proper MBC recall.

### ZFP318 confers recallability to otherwise poorly recallable MBCs

Given that ZFP318 is essential for MBC recallability, we wanted to test whether ZFP318 is sufficient for conferring proper recallability to otherwise poorly recallable MBCs. We created a transgenic system that allows inducible expression of ZFP318 in a B cell-specific manner. As shown in Fig. 4A, a single copy of TRE3G-ZFP318 expression cassette was knocked into the *Col1a1* safe locus to create *Col1a1*^Teton-ZFP318/+^ mice. These mice were further bred with *Rosa26*^LSL-rtTA3^;*Cd79a*^Cre/+^ mice. B cells from the resulting strain can be induced to express exogenous ZFP318, yielding a system to achieve <u>B</u> cell-<u>r</u>estricted induction of <u>s</u>ingle-copy <u>k</u>nocked-in ZFP318 gene expression. To simplify, we call this compounded transgenic strain BRISK^ZFP318^. We used *Col1a1*^Teton-ZFP318/+^;*Rosa26*^LSL-rtTA3/+^;*CD79a*^Cre/+^ (BRISK^ZFP318^) B1-8^hi^ mice or *Col1a1*^+/+^;*Rosa26*^LSL-rtTA3/+^;*CD79a*^Cre/+^ (BRISK^CTRL^) B1-8^hi^ mice as B cell donors in adoptive transfer experiments.

**Figure 4.**
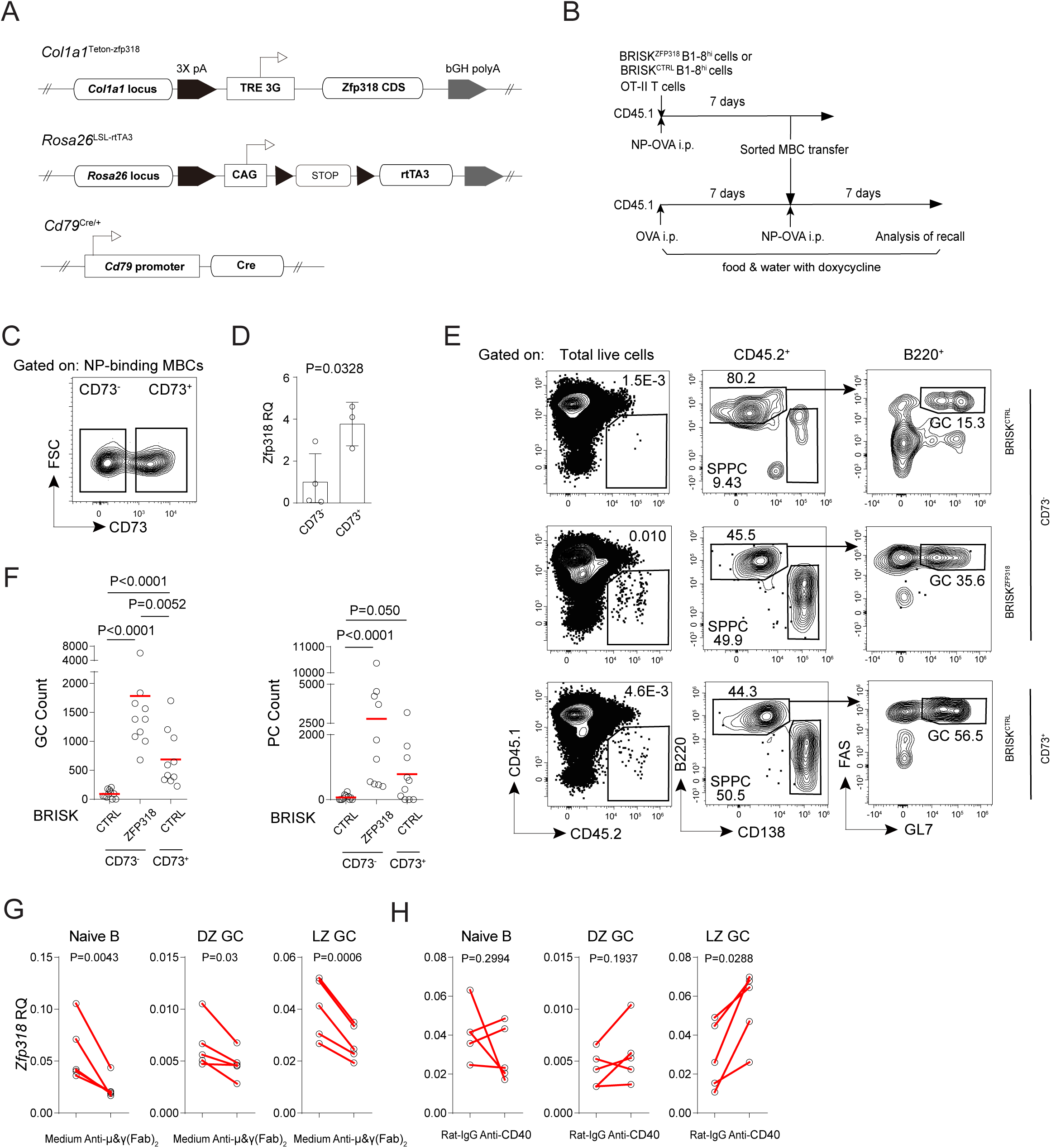
Enhanced recall by induced expression of exogenous ZFP318 and external stimuli that regulate ZFP318 expression. **(A)** Diagrams of *Col1a1*^Teton-zfp318^, *Rosa26*^LSL-rtTA3^ and *Cd79*^Cre/+^ alleles for the BRISK^ZFP318^ strain. **(B)** Experimental design. **(C)** The gating strategy to sort-purify CD73^+^ and CD73^-^ B1-8^hi^ MBCs. **(D)** Zfp318 expression in CD73^+^ and CD73^-^ B1-8^hi^ MBCs by qPCR. *P* values by Student’s *t* tests. **(E-F)** Gating strategy to identify donor-derived cells (**E**) and summary data (**F**) of GCs and splenic PCs derived from BRISK^CTRL^ CD73^-^, BRISK^ZFP318^ CD73^-^ and BRISK^CTRL^ CD73^+^ B1-8^hi^ MBCs 7 days after recall. Data were pooled from 2 independent experiments, with each symbol indicating one mouse and lines denoting means. *P* values by Mann-Whitney tests. **(G-H)** External stimuli that regulate ZFP318 expression. *Zfp318* mRNA levels in naïve B cells, DZ GC B cells, or LZ GC B cells after 4-hour stimulation with anti-IgM/IgG (**G**), anti-CD40 (**H**) or respective controls. Data were pooled from 5 independent experiments with each pair of circles indicating one experiment. *P* values by paired *t* tests.

To find a surface marker that can help enrich ZFP318-expressing MBCs, we tested CD73, which has previously been shown to enrich GC-derived MBCs^26,27^. As shown in Fig. S5, In NP-KLH-immunized AID-Cre;*Zfp318*^ZSDAT/+^ mice, ∼70% CD73^+^ NIP-binding MBCs were ZFP318-expressing, whereas the CD73^-^ MBC population contained ∼90% tdTomato^-^ MBCs. Therefore, CD73 is a useful marker to separate a population of 70%-pure ZFP318-expressing MBCs from a population of 90%-pure ZFP318-negative MBCs. To obtain sufficient numbers of sort-purified CD73^+^ and CD73^-^ B1-8^hi^ MBCs required for testing recall in lymphoreplete hosts, we harvested MBCs on day 7 post primary immunization. MBCs at this earlier time point are *bona fide* MBCs in that they are antigen-specific IgD^-^CD38^+^GL7^-^ cells, and they can respond to antigen re-challenge by producing secondary GCs and/or PCs. Moreover, they express similar transcriptomes as MBCs generated later and prominently contribute to the overall MBC pool (Ref. 28 and unpublished observation). We also verified that the recall function of day 7 B1-8^hi^ MBCs depended on ZFP318 (Fig. S4E-H), just as MBCs isolated on day 30.

Having established these parameters, we sort-purified CD73^-^ and CD73^+^ B1-8^hi^ MBCs of the BRISK^ZFP318^ background and CD73^+^ B1-8^hi^ MBCs of the BRISK^CTRL^ background, by the same protocol as described in Fig. 2H and Fig. S2B. These three groups of MBCs were transferred into three separate groups of CD45.1 mice that were primed with OVA and treated with doxycycline (Fig. 4B). At the time of cell transfer, sort-purified CD73^+^ NP-binding MBCs expressed much higher levels of ZFP318 mRNA than CD73^-^ counterparts (Fig. 4C-D), as expected. Following cell transfer, recipient mice were immunized with NP-OVA. As measured 7 days post immunization, CD73^-^ BRISK^ZFP318^ MBCs were functionally inferior to CD73^+^ counterparts, as expected; ZFP318 induction conferred a markedly improved recallability to CD73^-^ MBCs, leading to enhanced production of secondary GCs and PCs (Fig. 4E-F). Because these three groups of MBCs had essentially the same BCR, we can rule out affinity or isotype variation as the explanation for varied recall performance. We conclude ZFP318 itself is necessary and sufficient for conferring proper recallability to MBCs.

### ZFP318 expression is promoted by T-cell help

ZFP318 expression is increased in GC memory precursors. To identify potential factors that could induce ZFP318 upregulation, sort-purified light-zone or dark-zone GC B cells were treated with either anti-Ig to mimic antigen stimulation of BCR or agonistic anti-CD40 to mimic T-cell help. Naïve B cells were tested in parallel. Interestingly, whereas ZFP318 was downregulated by BCR stimulation in all types of B cells, it was upregulated by CD40 signaling and only in LZ GC B cells (Fig. 4G-H), suggesting that T-cell help plays a key role in promoting ZFP318 expression during affinity-based positive selection in the GC light zone.

### ZFP318 is required for mitochondrial integrity and for prevention of re-activation-induced cell death

Given the fact that ZFP318-deficient mice mount an intact primary response but an impaired recall response, the MBC recallability controlled by ZFP318 is likely a very upstream event in MBCs responding to antigen re-challenge, possibly involving a life-death decision. To explore this issue, we set up 3 systems to conduct pairwise transcriptomic comparisons of MBCs that differed in ZFP318 expression and recallabilities, with naïve B cells as additional controls (Fig. 5A-C). First, sort-purified EYFP^+^ NIP-binding MBCs from AID-Cre;*Zfp318*^ZSDAT/ZSDAT^;Rosa26-Ai3 mice (ZFP318-deficient) were compared to those from control AID-Cre;Rosa26-Ai3 mice (ZFP318-sufficient) 30 days after NP-KLH immunization (Fig. 5A; corresponding MBC recall experiment in Fig. 3C-E). Second, sort-purified tdTomato^+^ B1-8^hi^ MBCs of AID-Cre;*Zfp318*^ZSDAT/+^ mice (ZFP318-heterozygous) were compared to those of AID-Cre;*Zfp318*^ZSDAT/ZSDAT^ mice (ZFP318-deficient) 7 days after immunization (Fig. 5B; corresponding MBC recall experiment in Fig. S4E-H). Third, sort-purified tdTomato^+^ ZFP318-deficient B1-8^hi^ MBCs of AID-Cre;BRISK^ZFP3^^18^ (ZFP318-induced) or AID-cre;BRISK^CTRL^ (ZFP318-deficient) background 7 days after immunization under doxycycline treatment (Fig. 5C; AID-Cre instead of *CD79a*^Cre/+^ used here for the BRISK). The fact that induction of exogenous ZFP318 confers increased recallability was verified in Fig. 4.

**Figure 5.**
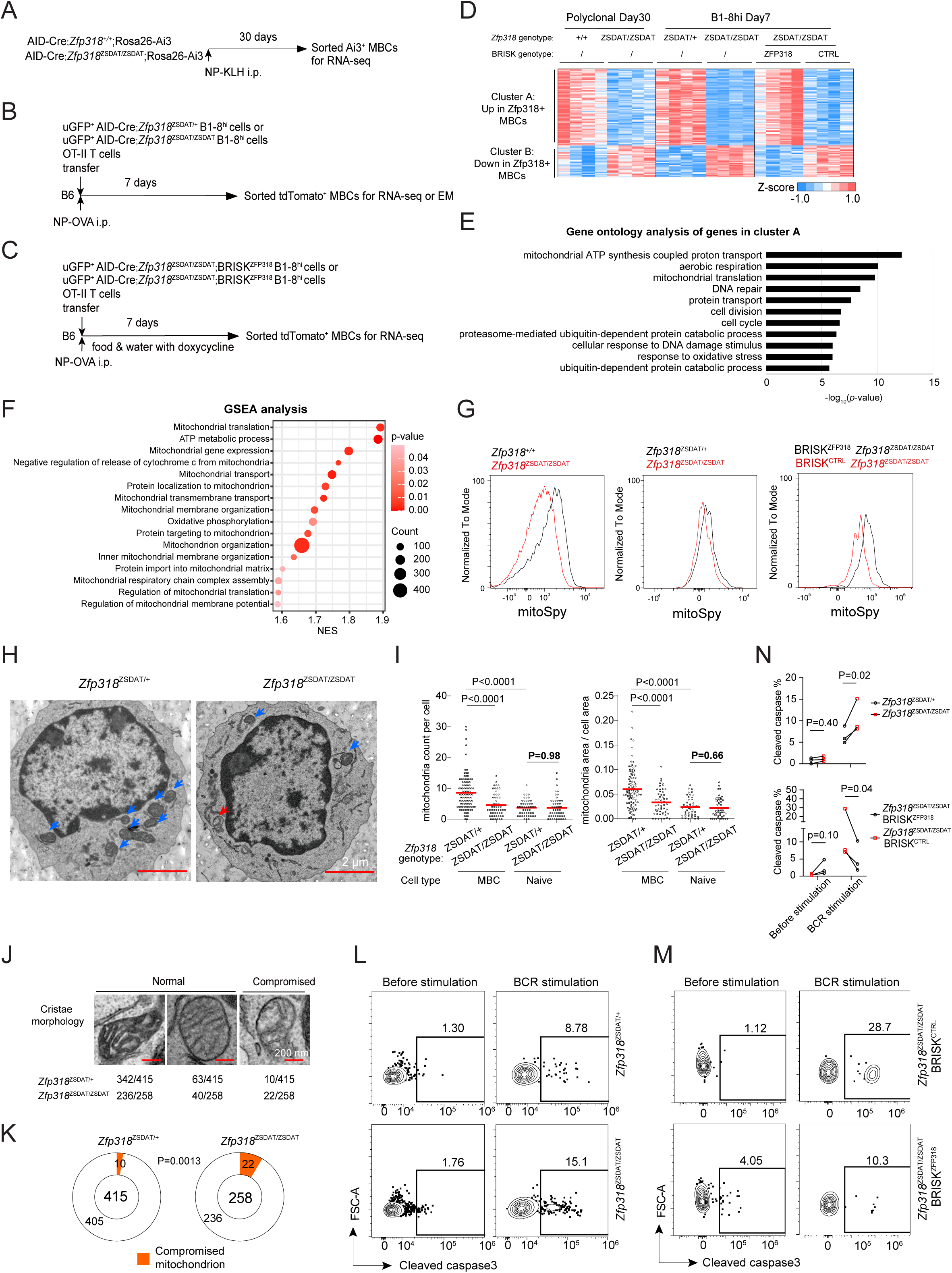
ZFP318 orchestrates a mitochondrial state conducive to recall. **(A-F)** Transcriptomic features of Zfp318-expressing (Zfp318^+^) MBCs compared to Zfp318-deficient (Zfp318^-^) MBCs. **(A-C)** Three experimental settings. note: AID-Cre instead of *CD79a*^Cre/+^ used for the BRISK here. **(D)** K-means clustering analysis of differentially expressed genes (*P*<0.05 at least in one experimental setting) pooled from the 3 experimental settings comparing Zfp318^+^ and Zfp318^-^ MBCs. **(E)** Gene ontology analysis of genes up-regulated in Zfp318^+^ compared to Zfp318^-^ MBCs (*P*<10^-5^). **(F)** GSEA analysis showing enrichment of mitochondrion-related pathways (*P*<0.05). Normalized enrichment score (NES), *p* values, and FDR are given in the graphs. **(G)** Histogram overlay of mitoSpy fluorescence of Zfp318^+^ and Zfp318^-^ MBCs in the 3 different experimental settings (**A-C**). **(H-K)** Electron microscopy of sort-purified B1-8^hi^ MBCs. **(H**) Representative micrographs of sort-purified B1-8^hi^ MBCs of indicated types. Blue arrow indicates normal mitochondrion, and red arrow indicates compromised one. (**I**) Nominal numbers (Left) and fractional areas (Right) of mitochondria in MBCs and naïve B cells (also see Fig. S6), quantitated from micrographs as in (**H**). Each symbol represent one cell, and lines denote means. Data were pooled from 2 independent experiments. *P* values by *t* tests. **(J)** Representative micrographs of cristae morphology in mitochondria. **(K)** Fractional abundance of structurally compromised mitochondria in tdTomato^+^ MBCs. *P* values by Fisher’s exact tests. **(L)** Representative profiles of cleaved caspase-3 expression by tdTomato^+^ B1-8^hi^ MBCs of an AID-Cre;*Zfp318*^ZSDAT/+^ or AID-Cre;*Zfp318*^ZSDAT/ZSDAT^ genotype, which were untreated or stimulated with an anti-IgM/G antibody (10 μg/ml) for 3 h. **(M)** Representative profiles of cleaved caspase-3 expression by tdTomato^+^ B1-8^hi^ MBCs of an AID-Cre;*Zfp318*^ZSDAT/ZSDAT^;BRISK^ZFP318^ or AID-Cre;*Zfp318*^ZSDAT/ZSDAT^; BRISK^CTRL^ genotype, which were untreated or stimulated with an anti-IgM/G antibody (10 μg/ml) for 3 h. **(N)** Summary data of percentage of cells containing cleaved (active) Caspase 3 in the indicated cell types before and after anti-IgM/G stimulation. Connected symbols are paired control and experimental groups from the same experiment. *P*-values by ratio paired *t* tests.

By K-means clustering analyses, we observed a total of 12 gene clusters that were differentially expressed in these three datasets (Fig. S6A). The cluster A of 812 genes showed consistent downregulation, and the cluster B of 343 genes showed consistent upregulation in ZFP318-deficient MBCs as compared to ZFP318-sufficient, ZFP318-heterozygous or ZFP318-induced MBCs (Fig. 5D). By gene ontology analysis, the cluster A was highly enriched with mitochondrion-related, nuclear genome-encoded genes, including those that support mitochondrial ATP synthesis-coupled proton transport and mitochondrial translation and those mediating cellular responses to DNA damage and oxidative stress (Fig. 5E). Gene set enrichment analysis further showed, in ZFP318-deficient MBCs, downregulation of 16 mitochondrion-related gene sets that impinge on all aspects of mitochondrial structure and function (Fig. 5F). In particular, ZFP318-deficient MBCs exhibited markedly reduced expression of genes orchestrating mitochondrial translation (Fig. S6B), ATP synthesis (Fig. S6C), defense against oxidative stress (Fig. S6D-F) and negative regulation of cytochrome c release (Fig. S6G). On the other hand, none of these differences were observed when naïve B cells of these genotypes were compared.

Consistent with those transcriptomic features, mitochondria of ZFP318-deficient MBCs are structurally and functionally compromised. By MitoSpy staining, ZFP318-deficient MBCs consistently exhibited lower mitochondrial membrane potentials as compared to ZFP318-sufficient, ZFP318-heterozygous and ZFP318-induced counterparts (Fig. 5G, quantitation in Fig. S6H). Compared to ZFP318-sufficient MBCs, ZFP318-deficient MBCs contained fewer mitochondria (Fig. 5H-I). In ZFP318-deficient MBCs, a significantly larger fraction of mitochondria showed a swollen morphology with few intact cristae (Fig. 5J-K), a type of structural deformation that often results from abnormally increased mitochondrial permeabilities. On the other hand, mitochondria in ZFP318-deficient naïve B cells did not show any numerical reduction or structural compromise (Fig. S6I and Fig. 5I).

BCR activation significantly increases mitochondrial activities and causes ROS-related mitochondrial stress, which could lead to B cell death^29^. The ZFP318 deficiency may cause MBCs to be more sensitive to antigen reactivation-induced cell death. Consistent with this possibility, brief anti-Ig stimulation led to a significantly more pronounced increase in Caspase 3 activation in Zfp318-deficient MBCs, whereas induction of exogenous ZFP318 in ZFP318-deficient MBCs suppressed this tendency to die of reactivation (Fig. 5L-N). Together, these data strongly suggest that ZFP318 promotes recallability by counteracting reactivation-induced cell death that stems from excessive ROS and oxidative stress.

### An ROS scavenger rescues the recall defect of ZFP318-deficient MBCs

To test whether oxidative stress contributes to the poor recall performance of ZFP318-deficient MBCs, MBCs were enriched from NP-KLH-immunized wildtype or ZFP318-deficient mice as T/GC/PC-depleted CD19^+^ cells by essentially the same protocol as in Fig. 3C. We transferred equal numbers of ∼2000 NIP-binding MBCs of the two genotypes into separate OVA-primed CD45.1 recipients, which were then immunized with NP-OVA (Fig. 6A). Immunized mice were intraperitoneally given ROS scavenger NAC or PBS every the other day. As shown in Fig. 6B-C, whereas NAC treatment did not change the recall response of wildtype MBCs, it significantly improved the recall generation of secondary GCs and PCs by ZFP318-deficient MBCs. The endogenous primary response by CD45.1 host cells was comparable between NAC and PBS groups (Fig.S5C). Together, these data rule out the possibility that ZFP318-deficient MBCs would home to unknown tissue niche without access to immunizing antigen or they simply die off upon adoptive transfer before antigen reactivation. Instead, they indicate that ZFP318-controlled recallability rests in ZFP318-mediated protection of MBCs from oxidative stress, which would otherwise cause MBCs to die while being reactivated by antigen.

**Figure 6.**
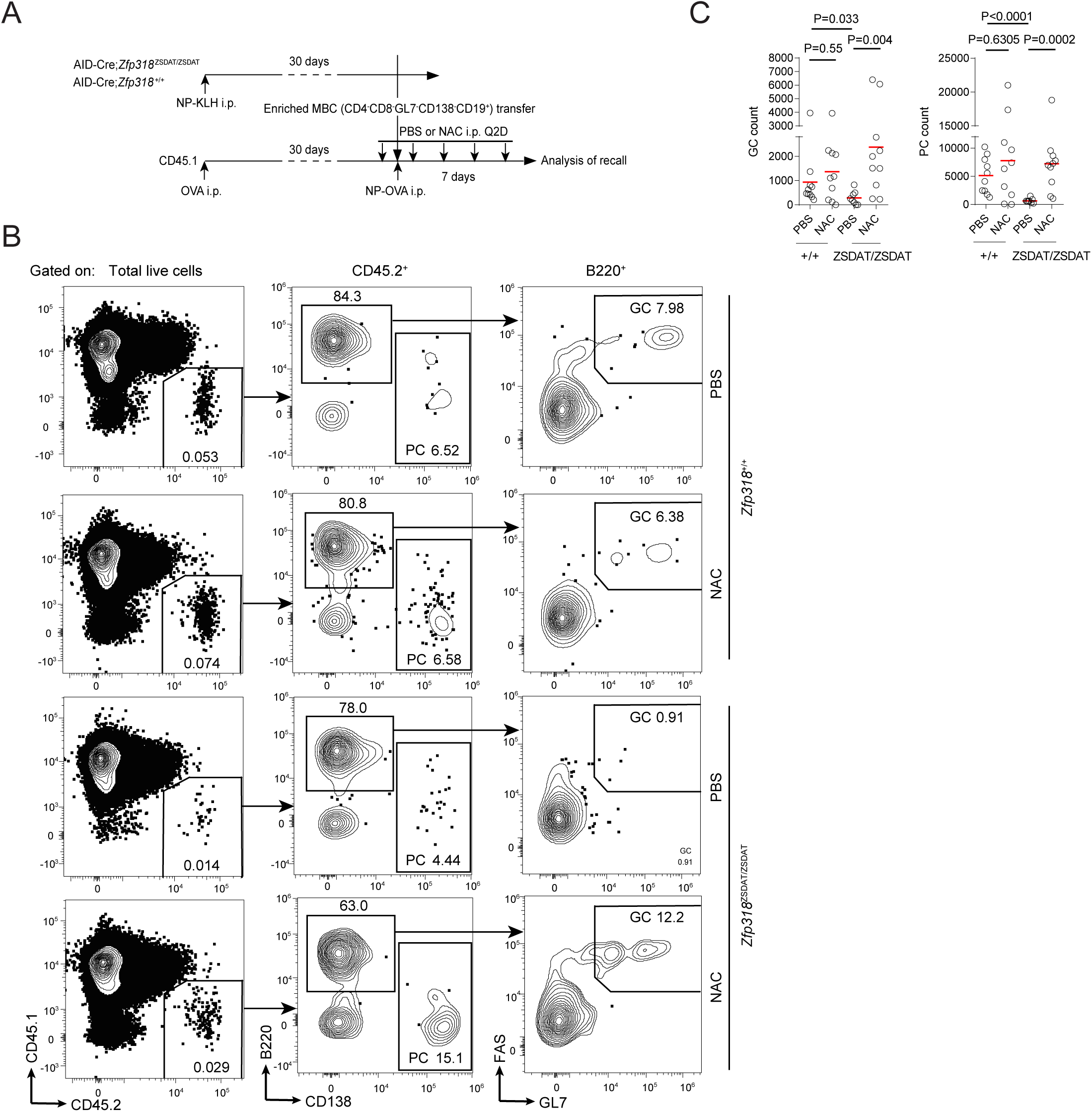
ROS scavenger rescues recall defects in ZFP318-deficient MBCs. **(A)** Experimental design. **(B)** Gating strategies to identify CD45.2^+^ GCs and SPPCs. **(C)** Summary data of cell abundance of indicated types. Data were pooled from 2 independent experiments, with each symbol indicating one mouse and lines denoting means. *P* values by Mann-Whitney tests.

### Re-programming of poorly recallable MBCs by exogenous ZFP318

MBCs that do not express ZFP318 are enriched with MBCs developed from a GC-independent route. These MBCs may harbor clones that react to antigenic variants of pathogens ^30^, a feature that could be exploited for development of broadly neutralizing vaccines. To establish a proof-of-principle method for re-programing poorly recallable GC-independent MBCs, we resorted to mRNA transfection of CD73^-^ MBCs, which are mainly GC-independent cells. Polyclonal MBCs resulting from primary immunization were transfected with lipid nanoparticles (LNP) containing ZFP318- or GFP-coding mRNA and tested for recall after adoptive transfer (Fig. 7A). As shown in Fig. 7B-C, although CD73^-^ MBCs transfected with LNP-GFP were not recalled as efficiently as the CD73^+^ counterpart to produce GCs or PCs, LNP-ZFP318 transfection of CD73^-^ MBCs led to a much stronger recall response generating both GCs and PCs.

**Figure 7.**
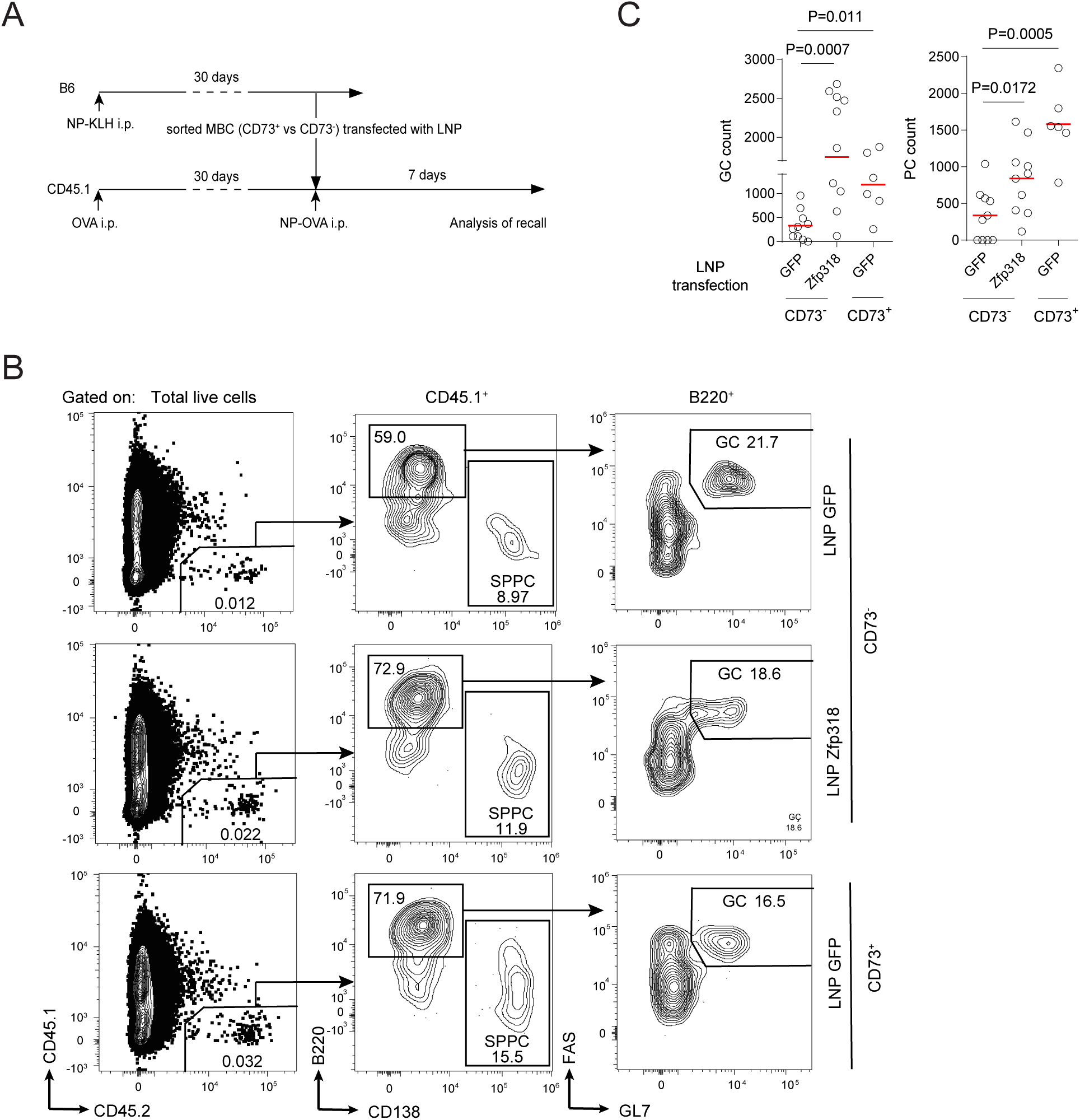
Reprogramming recallability by exogenous ZFP318. **(A)** Recall experiments using sorted CD73^+^ or CD73^-^ MBCs that were transfected with LNP carrying control GFP or Zfp318 mRNAs. **(B-C)** Gating strategy to identify donor-derived cells (**B**) and summary data (**C**) of GCs and splenic PCs derived from control CD73^-^, Zfp318-induced CD73^-^ and control CD73^+^ MBCs 7 days after recall. Data were pooled from 2 independent experiments, with each symbol indicating one mouse and lines denoting means. *P* values by Mann-Whitney tests.

### Abundance of ZFP318-expressing MBCs predicts the quality of vaccine-induced antibody responses

Given the central role of ZFP318 in controlling the MBC recallability, strengths of antibody immunity induced by prime-boost vaccination would be predicted by the abundance of ZFP318-expressing MBCs rather than the abundance of total MBCs. We validated this idea in three published systems of prime-boost vaccination, in each of which two related but distinct vaccine formulations have been compared and shown to have different efficacies. First, a colloidal manganese salt (Mn jelly, MnJ) is demonstrably a superior adjuvant that, in part by activating cGAS-STING and NLRP3 pathways, stimulates much stronger antibody responses as compared to what is achievable with alum as the alternative adjuvant ^31,32^. As shown in Fig. S7A-B, MnJ-adjuvanted immunization induced more ZFP318-expressing MBCs after the prime, whereas total antigen-binding MBCs were comparable to what was achieved with alum. Second, a tripartite fusion of the receptor-binding domain (RBD) of SARS-CoV2 S protein, type I interferon and immunoglobulin Fc domain (I-R-F) has been shown to be a superior vaccine candidate than alum-adjuvanted RBD, as demonstrated by much higher post-boost titers of total and neutralizing anti-RBD antibodies ^33^. As shown in Fig. S7C-D, the I-R-F regimen induced more ZFP318-expressing RBD-specific MBCs as measured right before the boost, whereas total RBD-binding MBCs were comparable between the two groups. Finally, we have previously compared two formulas of mRNA vaccines targeting the SARS-CoV2 RBD, one expressing the RBD dimer and the other expressing trimerized RBD (3×RBD). The two formulas induced a very similar primary response in terms of GCs, PCs, total RBD-binding MBCs, and RBD-binding antibody titers, as measured right before the boost. After the boost, however, 3×RBD immunization gave rise to markedly higher titers of serum RBD-specific antibodies that persisted long after the boost ^34^. As shown in Fig. S7E-F, the abundance of ZFP318-expressing but not total RBD-binding MBCs, as measured before the boost, was much higher with 3×RBD than with RBD immunization. Taken together, in three independent scenarios of prime-boost vaccines, the better post-boost performance is correlated with ZFP318-expressing MBCs but not total antigen-binding MBCs. These data are consistent with the fact that ZFP318-expressing MBCs are the functionally dominant B cell memory compartment and ZFP318 is a master regulator of MBC recallability.

## Discussion

By fate-mapping, cell deletion, gene ablation, inducible complementation experiments, we show that all MBCs are not equally recallable, and functional memory recall is predominantly carried out by ZFP318-expressing MBCs. ZFP318 is dispensable for naïve B cell activation but essential for MBCs to properly respond to antigen and produce secondary GCs and PCs. ZFP318 controls genes that mediate vital mitochondrial functions, particularly those responsible for oxidant detoxification, which appear to be particularly important for MBCs. Given the stark contrast between ZFP318-deficient MBCs and naïve B cells, future studies are necessary to elucidate how ZFP318 controls those mitochondrion-related genes and identify co-factor(s) that contextualizes the cell type-specific gene regulation by ZFP318. Given the fact that exaggerated oxidative stress could induce ferroptosis and the fact that GPX4, a gene subjected to ZFP318-dependent regulation, prevents ferroptosis ^35^, it would be interesting to explore whether and how ferroptosis may be involved in MBC biology and regulated by ZFP318.

Antigen-specific MBCs develop from both GC-independent and -dependent routes^5,26,36–38^. with the former affording potentially a broader spectrum of reactivities toward the antigen and the latter providing a more focused, affinity-matured response. GC-independent MBCs contain mainly unswitched IgM^+^ cells together with a minor component of IgG^+^ MBCs, whereas GC-dependent MBCs are composed of both switched and IgM^+^ MBCs^26,38^. Given the fact that ZFP318-expressing MBCs are enriched with GC-derived cells, whereas MBCs without ZFP318 expression are enriched with GC-independent ones, our data suggest a major difference in recallability between MBCs derived from the two routes. Previous human studies suggest that long-lived and functionally recallable MBCs are primarily those with GC experience^39–41^, although mouse work shows that GC-independent MBCs can be recalled at least in lymphopenic hosts^38^. It is important to point out that antigen-specific MBCs in the current work by definition do not express IgD. Some antigen-activated B cells can become quiescent MBCs without losing IgD expression^42^. We do not yet know whether such IgD^+^ MBCs, mostly GC-independent ones, maintain ZFP318 expression and how they would perform in recall.

The concept of recallability differ from the MBC recall fate bifurcation, or, the ability of reactivated MBCs to make choice between secondary GCs and PCs^1^. Recallability is regulated at a more fundamental level than the fate bifurcation in that MBCs must be able to survive and productively respond to antigen before realizing their fate choice. Indeed, the absence of ZFP318 always affects recall production of both secondary GCs and PCs but never one over the other. Importantly, the intrinsic recallability of MBCs can be most rigorously assessed by the recall assay after adoptive transfer, even though such assay for polyclonal MBCs in lymphoreplete animals is technically challenging and not routinely practiced in the field. While functionally dominant, ZFP318-expressing MBCs represent a numerical minority in total antigen-binding MBCs. ZFP318-expressing MBCs do not co-segregate with any MBC subsets classified according to isotypes or CD80 and PDL2 surface markers. In fact, the IgM^+^ MBC subset or the CD80^-^PDL2^-^ MBC subset each contains only 10% or fewer ZFP318-expressing cells, indicating that most of them are poorly recallable and their observable preponderance for development into secondary GCs is carried out by a minority within.

Given the fact that all MBCs are not equally recallable, it is an intriguing question as to why and how the immune system could make so many MBCs with so little ZFP318 expression. Our results indicate that T cell help signals such as CD40L promote ZFP318 expression in GC light-zone cells. ZFP318-expressing MBCs may represent cells that have experienced sufficient amounts of T cell help to be considered “licensed” clones, allowed to efficiently re-participate in the secondary response. By the same token, the large contingency of MBCs with little or no ZFP318 expression could result from limited access to T cell help during development. This idea is consistent with the fact that MBCs expressing no ZFP318 are enriched with GC-independent cells. Moreover, in our B1-8^hi^/OT-II co-transfer model, the proportion of ZFP318-expressing MBCs in total antigen-specific MBCs appeared to increase as more transferred OT-II cells were provided (unpublished observation). Our finding is also consistent with the observation that a majority of MBC clones do not re-participate in secondary GCs following counter-lateral boost, whereas those clones that do re-participate are those of high germline affinities and presumably having better chances of receiving more T cell help^43^. MBCs with little or no ZFP318 expression may represent clones of unproven safety for participation in the secondary response but at the same time may serve as a repertoire of broader reactivities, only to be productively recalled when exuberant T cell help or additional survival factors such as BAFF becomes abundantly available^44,45^.

From a perspective of prime-boost vaccines, ZFP318-expressing MBCs could serve as a good biomarker for stronger and more durable boost performance. Treatments to induce ZFP318 expression in otherwise non-expressing MBCs, as proved by concept with mRNA transfection experiments, could be a useful strategy to increase the vaccine potency and to adjust the breadth of vaccine-induced antibody responses.

### Limitation of the Study

How expression of ZFP318 is regulated remain to be further clarified. For example, how CD40 and BCR signaling antagonistically regulates ZFP318 expression and whether and how the level of ZFP318 expression in individual MBCs is set according to their histories of interactions with T cells are but two intriguing questions yet to be answered. The current study has also not elucidate how ZFP318 controls, directly or indirectly, those mitochondrial genes responsible for oxidant detoxification. It remains to be confirmed whether ZNF318, the human homolog of ZFP318, plays a similar role in regulating MBC biology.

## Acknowledgements

We thank Dr. M. Nussenzweig for providing the B1-8^hi^ mice. The work was funded in part by the National Key R&D Program of China (Ministry of Science and Technology, 2018YFE0200300 to H.Q.), National Natural Science Foundation of China (grant 31830023, 81621002 to H.Q., grant 31900629 to Y.W. and grant 32200725 to W.S.), China Postdoctoral Science Foundation (grant 2022T150351 to W.S.), the Changping Laboratory, the Tsinghua-Peking Center for Life Sciences, the Beijing Municipal Science & Technology Commission, and the Beijing Frontier Research Center for Biological Structure. H.Q. is a New Cornerstone Investigator.

## Author contributions

H.Q. conceptualized and supervised the study. Y.W. and W. Shao conducted a majority of the experiments. W. Shao conducted RNA-seq analyses. X.L. made the initial design of the ZSDAT reporter. J.L. and G.Y. designed and prepared LNP. W. Shi conducted ELISA. Q.L. helped with mRNA vaccine experiments under the supervision of L.Z. H.Q. and Wen S. wrote the paper with input from all authors.

## Competing interest

Y.W. and H.Q. are co-founders of Emergent Biomed Solutions, Ltd.

## Data availability

All data generated during and/or analyzed during the current study are available from the corresponding author upon reasonable request. Source data are included in this published article (and its supplementary information files).

## STAR Methods

### RESOURCE AVAILABILITY

#### Lead contact

Further information and requests for resources and reagents should be directed and will be fulfilled by the lead contact Hai Qi.

#### Materials availability

Mouse lines generated in this study are subject to MTA agreements.

#### Data and code availability

RNA-seq data have been deposited at GEO and are publicly available as of the date of publication. Accession numbers are listed in the key resources table. Any additional information reported in this paper will be shared by the lead contact upon request.

### EXPERIMENTAL MODEL AND STUDY PARTICIPANT DETAILS

#### Animal experimental models

C57BL/6 (Jax 664), GFP-transgenic (Jax 4353), AID-Cre (Jax 018422), Rosa26-Ai3 (Jax 7903), OVA_323–339_-specific T-cell receptor transgenic OT-II (Jax 4194), Rosa26-CAGs-rtTA3 (Jax 029627), and CD45.1 (Jax 2014) mice were originally from the Jackson Laboratory. The B1-8^hi^ Ig heavy chain knock-in mice were a kind gift of Dr. M. Nussenzweig. *Cd79a*^cre/+^ mice were a gift from Dr. M. Reth. The ZFP318-reporting *Zfp318*^ZSDAT/+^ knock-in/knock-out strain mice and ZFP318-inducing *Col1a1*^Teton-ZFP318/+^ knock-in strain mice were developed at BIOCYTOGEN by CRISPR/Cas9 technology. Ighd KO mice were developed at GemPharmatech by CRISPR/Cas9 technology. Relevant alleles were interbred to obtain desired genotypes. All mice were maintained under specific pathogen-free conditions, and used in accordance of governmental and Tsinghua guidelines for animal welfare.

### METHOD DETAILS

#### Immunization and viral infection

For immunization with SRBC, 10^8^ SRBC in 1 ml suspension (Beijing Solarbio Science & Technology) was intraperitoneally injected to each mouse. To measure the NP-specific response in various experiments, mice were intraperitoneally injected with 100 μg NP-KLH, 100 μg or 20 μg NP-OVA (Biosearch Technologies, 20 μg for recall) mixed with 2 μg LPS (Sigma) in alum (Thermo Scientific). To prepare carrier-primed mice for MBC transfer, mice were intraperitoneally immunized with 20 μg OVA mixed with 2 μg LPS in alum. To compare adjuvants, mice were intraperitoneally immunized with 100 μg NP-KLH mixed with lipopolysaccharide in alum or mixed with 300 μg MnJ (MnStarter Biotechnology), as previously described adjuvant^31,32^. To compare RBD and 3×RBD trimer mRNA vaccines, mice were intramuscularly immunized with 10 μg respective mRNA LNP. To compare recombinant RBD dimer or IFN-Pan-RBD-Fc vaccines, mice were immunized with 3.4 μg RBD dimer or 10.4 μg IFN-Pan-RBD-Fc protein, as previously described^33^.

#### Flow cytometry

To profile different cell populations by flow cytometry, single-cell suspension was washed with PBS, and then stained with indicated antibodies in MACS buffer (PBS supplemented with 1% FBS and 5 mM EDTA). Staining reagents included FITC anti-IgG3, streptavidin-PE and BV421 Goat anti-Rabbit IgG from BD Biosciences; AF700 anti-CD38, EF450 anti-GL7, APC anti-CD80, FITC anti-PD-L2, APC anti-CD45.1 and FITC anti-CD45.2 from Thermo Fisher Scientific; PerCP Cy5.5 anti-IgD, PE-Cy7 anti-CD95, allophycocyanin (APC)-Cy7 anti-B220, BV510 anti-CD138, FITC anti-IgG1, FITC anti-IgG2a, FITC anti-IgG2b, BV421 anti-IgM, PerCP Cy5.5 anti-Gr1, PE-CD73, streptavidin-APC, anti-Ig light chain λ and MitoSpy™ NIR DiIC1(5) from Biolegend; FITC anti-IgG2c from Bethyl. AF700 anti-CD38, Isotype-matched control antibodies were also purchased from these respective vendors. Surface staining was done on ice with primary reagents incubated for 30 minutes followed by secondary reagents for 30 minutes with washes in-between. To detect NP-binding cells in GC B cells, MBCs, SPPCs or BMPCs, NP_(38)_- or NP_(44)_-PE or biotinylated NIP_(15)_-BSA (Biosearch Technologies) were used. For caspase detection, cells were firstly stained with surface markers and then were fixed and permeabilized by Cytoperm/Cytofix kit (BD Biosciences) following manufacturer’s manuals. Cells were then stained with primary antibody of cleaved caspase 3 (Cell Signaling Technology), followed by staining with secondary antibody BV421 anti-Rabbit IgG (BD Biosciences). All cytometric data were collected on an LSR II, or an FACSAria III cytometer (BD Biosciences) or an Aurora cytometer (Cytek Biosciences) and analyzed with the FlowJo software (TreeStar). Dead cells and non-singlet events were excluded from analyses based on staining with Zombie Yellow (Biolegend) and characteristics of forward and side scatters.

#### Recall assays of MBCs

First, to measure the recall response by polyclonal MBCs in a recipient, AID-Cre;*Zfp318*^+/ZSDAT^ mice were immunized and used as the donor of MBCs 30 days after primary NP-KLH immunization. To isolate MBCs, donor splenocytes were depleted of CD4^+^, CD8^+^ cells by CD4 and CD8 Microbeads (Miltenyi Biotec), and GL7^+^, CD138^+^, IgD^+^ cells using a cocktail of biotinylated anti-GL7, anti-CD138, anti-IgD antibodies (Biolegend) in combination of streptavidin Microbeads (Miltenyi Biotec). CD19^+^ cells were then enriched using CD19 Microbeads (Miltenyi Biotec). The resulting cell preparation was FACS-analyzed, normalized for the abundance of NIP-binding B220^+^CD138^-^GL7^-^FAS^-^CD38^+^IgD^-^ MBCs, and transferred into CD45.1 recipients that were primed with carrier protein OVA 30 days before. After cell transfer, recipient mice were immunized with NP-OVA. To delete ZFP318-expressing cells, these mice were given 5 μg/kg diphtheria toxin (Sigma) or PBS intraperitoneally. The recall response by CD45.2^+^ donor-derived MBCs was measured 7 days after NP-OVA immunization. Secondly, to measure the recall response by MBCs carrying NP-specific BCRs in a secondary host, AID-Cre;*Zfp318*^+/ZSDAT^ GFP-expressing B1-8^hi^ cells were adoptively transferred into B6 mice that also received OT-II T cells and were immunized with NP-OVA. Resulting NP-binding B220^+^CD138^-^ GL7^-^FAS^-^CD38^+^IgD^-^GFP^+^ MBCs from these primary hosts were sort-purified into tdTomato^+^ and tdTomato^-^ cells, and these latter cells were transferred into separate groups of secondary B6 recipients that were primed with OVA protein 30 days before. These recipient mice were then immunized with NP-OVA, the recall response by GFP^+^ B1-8^hi^ MBCs was measured 7 days after NP-OVA immunization. Third, to measure the recall response by ZFP318-deficient MBCs, AID-Cre;*Zfp318*^ZSDAT/ZSDAT^;Rosa26-Ai3 or control AID-Cre;*Zfp318*^+/+^;Rosa26-Ai3 mice were immunized and used as the donor of MBCs 30 days after primary NP-KLH immunization. To isolate MBCs, donor splenocytes were depleted of CD4^+^, CD8^+^ cells by CD4 and CD8 Microbeads, and GL7^+^, CD138^+^ cells using a cocktail of biotinylated anti-GL7 and anti-CD138 antibodies in combination of streptavidin Microbead. CD19^+^ cells were then enriched using CD19 Microbeads. The resulting memory B cell-enriched cell preparation was FACS-analyzed, normalized for the abundance of NIP-binding B220^+^CD138^-^GL7^-^ FAS^-^CD38^+^ MBCs, and transferred into CD45.1 recipients that were primed with carrier protein OVA 30 days before. After cell transfer, these recipient mice were immunized with NP-OVA. The recall response by donor-derived MBCs was measured 7 days after NP-OVA immunization. Fourth, to measure the recall response by ZFP318-deficient MBCs carrying NP-specific BCRs in a recipient, uGFP-expressing AID-Cre;*Zfp318*^+/ZSDAT^ B1-8^hi^ cells or AID-Cre;*Zfp318*^ZSDAT/ZSDAT^ B1-8^hi^ cells were adoptively transferred into B6 mice that also received OT-II T cells and were immunized with NP-OVA. Resulting NP-binding B220^+^CD138^-^GL7^-^FAS^-^ CD38^+^GFP^+^tdTomato^+^ MBCs from these primary hosts were sort-purified, and these cells were transferred into separate groups of secondary B6 recipients that were primed with OVA protein 30 days before. These recipient mice were then immunized with NP-OVA, the recall response by GFP^+^ B1-8^hi^ MBCs was measured 7 days after NP-OVA immunization. Fifth, to measure the recall response by MBCs with induced Zfp318 expression, BRISK^ZFP318^ or BRISK^CTRL^ GFP-expressing B1-8^hi^ cells were adoptively transferred into B6 mice that also received OT-II T cells and were immunized with NP-OVA. Resulting NP-binding B220^+^CD138^-^GL7^-^FAS^-^CD38^+^IgD^-^GFP^+^ MBCs from these primary hosts were sort-purified into CD73^+^ and CD73^-^ cells, and these latter cells were transferred into separate groups of secondary B6 recipients that were primed with OVA protein and fed with doxycycline (Beyotime). These recipient mice were then immunized with NP-OVA, the recall response by GFP^+^ B1-8^hi^ MBCs was measured 7 days after NP-OVA immunization. For recalls with LNP, sorted MBCs were transfected with LNP carrying either Zfp318 or GFP mRNAs (1 μg RNA/10^5^ cells) and incubated in RPMI at 37°C for 3 hrs. Cells were then washed by PBS, and prepared for adoptive transfer. For recalls with NAC treatment, recipient mice were given 150 mg/kg of acetylcysteine (MCE) or PBS intraperitoneally every the other day as indicated.

#### Encapsulation of RNA by LNP

LNP was fabricated by extruding two phases (aqueous phase and ethanol phase). Ethanol phase: The cationic lipid, cholesterol, PEG-lipid, and DSPE were used as the lipid phase to fabricate LNP. The four components were dissolved into pure ethanol. To facilitate dissolution, the solution was slightly heated by a heat gun (hair dryer) before using. DSPE and PEG-lipid would separate from ethanol solution at 4 ℃. The concentration of the cationic lipid, cholesterol, PEG-lipid, and DSPE is 43 mg/ml, 20 mg/ml, 10 mg/ml, and 9 mg/ml, respectively. The volume of cationic lipid, cholesterol, PEG-lipid, and DSPE is 100 μl, 100 μl, 50 μl, and 100 μl, respectively. After adding the four components together, 100 μl ethanol was added. The final volume of ethanol was 450 μl. Aqueous phase: RNA was dissolved into 50 mM citrate buffer with a concentration of 100 μg/900 μl. After loading the two phases into corresponding syringes, the two syringes were extruded with the speed ratio 1:3 (ethanol phase: buffer phase). The extruded LNP was dialyzed against PBS for 2 hours and water for 3 hours. After the dialyzation process, to check the quality of LNP, the diameter, polydispersity index, zeta potential of LNP were tested by Zetasizer Nano (Malvern Instruments, Malvern, UK) at least 5 times. All experiments were carried out using freshly packaged LNP within 12 h.

#### V_H_186.2 sequence analysis of NP-specific B cells

NIP-specific MBCs were identified by staining with biotinylated NIP-BSA (Biosearch Technologies) at 330 ng/ml for 30 min on ice and then with streptavidin APC. NIP-binding GL7^-^FAS^-^B220^+^CD138^-^IgD^-^ CD38^+^ MBCs were sort-purified 14 or 28 days after NP-KLH immunization. After incubation at 60 °C for 5 min in the lysis buffer, lysates of multiple 200-cell sorts per experiment was subjected to reverse-transcription by with the Superscript cDNA Synthesis Kit (Invitrogen) using the manufacturer’s suggested protocol. V_H_186.2 fragments were amplified from the complementary DNA by nested PCR using the following primers (first and second sense: 5′-TTCTTGGCAGCAACAGCTACA, first antisense including Cγ1 antisense: 5′-GGATCCAGAGTTCCAGGTCACT and Cμ antisense 5′-AAATGGTGCTGGGCAGGAAG; second antisense including Cγ1 antisense: 5′-GGAGTTAGTTTGGGCAGCAG and Cμ antisense 5′-AGCCCATGGCCACCAGATT). PCR products were purified by gel electrophoresis and cloned into a T vector (Takara), and individual bacterial colonies were picked for sequencing. Identical V_H_ sequences were counted only once as one clone, and the final result was compiled with unique clones for each category.

#### Quantitative RT-PCR

Quantitative RT-PCR was performed with a SYBR *Premix Ex Taq* II (Tli RNaseH Plus) (RR820; Takara) on a 7500 Real-Time PCR system (Thermo Fisher) or CFX Connect™ Real-Time PCR system (Bio-Rad). Primers used were: *Actb* sense 5′-GCTTCTTTGCAGCTCCTT, antisense 5′-TGCCAGATCTTCTCCATGT; *Zfp318* sense 5′-ACGTTCTCCTGGTCTGTGTTC, antisense 5′-AGTGGTCATTGCCAACTGTG. *Dtr* sense 5′-AGGGAAGAAGAGGGACCCAT, antisense 5′-AACATGAGAAGCCCCACGAT.

#### Measurement of serum NP-specific IgG by ELISA

Sera were harvested from relevant mice and stored at −80°C until use. 96-well ELISA plates (42592, Costar) were coated with NP_2_-BSA or NP_30_-BSA (Biosearch Technologies) in PBS overnight and washed. A serial 1:2 dilution of each serum sample, starting from 1:5000, was then loaded into the plate and incubated at 37 °C for 1 h. After washings, the plate was further incubated with HRP-conjugated goat anti-mouse IgG (Biolegend) at 1:6,000 for 1 h before development with TMB Substrate Set (Biolegend). The chromogenic reaction was stopped with 1 M HCL, and optical density (OD) was read on an iMark plate reader (Bio-Rad). For each group of animals, the resulting dilution curves were fit with the 3-parameter dose–response curve in Prism (Graphpad). Curves from two groups in comparison were analyzed by the extra sum-of-squares F-test, and their nominal EC50s were used to calculate fold-change in NP-specific IgG titers.

#### Electron Microscopy

Sort-purified MBCs or naïve B cells were fixed with 2.5% glutaraldehyde in 0.1M PB buffer for 2 h and then rinsed 3× 10 min with the PB buffer. Post-fixation staining was performed with 1% osmium tetroxide/K_3_Fe(CN)_6_ for 30 min. After washing in distilled water 3× 5 min, cells were dehydrated in a graded ethanol series (50%, 70%, 80%, 90%, 100%, 100%) for 5 min each. After the final incubation in 100% ethanol at the room temperature, cells were incubated in a 1:1 mix of ethanol and acetone for 5 min and then in 100% acetone for 5 min. Cells were then infiltrated and embedded in Pon 812 resin. The blocks were trimmed, and 70-nm sections were contrast-stained with uranyl acetate and citrate. Cells were then examined with TEM (Hitachi, H7650). Numbers of mitochondria per cell section, fractional mitochondrion-occupied area, and mitochondria density were quantitated with ImageJ.

#### RNA-seq and data analysis

Approximately 200-1000 cells of desired type were lysed in 6 μl lysis buffer (0.2% Triton X-100 and 2 U/μl of RNase inhibitor). Lysates were mixed with 3 μl of anchored oligo-dT primer (10 μM, 5′-AAGCAGTGGTATCAACGCAGAGTACT30VN-3′) and 3 μl of dNTP mix (10 mM, Fermentas), denatured at 72 °C for 3 min and immediately placed on ice afterwards. The first-strand reaction mix containing 1 μl SuperScript II reverse transcriptase (Invitrogen), 0.5 μl RNAse inhibitor (Clontech), 4 μl Superscript II First-Strand Buffer (Invitrogen), 1 μl DTT (100 mM, Invitrogen), 4 μl betaine (5 M, Sigma), 0.12 μl MgCl_2_ (1 M, Sigma), 0.2 μl TSO (100 μM) and 0.58 μl nuclease-free water, were added to each sample. Reverse transcription reaction was carried out by incubating at 42 °C for 90 min, followed by 10 cycles of (50 °C for 2 min, 42 °C for 2 min). Finally, the reverse transcriptase was inactivated by incubation at 70 °C for 15 min. PCR mix containing 25 μl KAPA HiFi HotStart ReadyMix, 0.5 μl IS PCR primers (10 μM) and 1.1 μl nuclease-free water were added to each sample. The reaction was incubated at 98 °C for 3 min, then cycled 17 times (98 °C 20 s, 67 °C 15 s, 72 °C 6 min), with a final extension at 72 °C for 5 min. cDNAs were purified by Vazyme DNA clean beads following manufacturer’s instructions. 1 ng of cDNA were used for the tagmentation reaction using Tn5 (Vazyme). The tagmentation reaction (20 μl) was incubated at 55 °C for 10 min and stopped by addition of 2 μl 0.2% SDS and incubated at 70 °C for 20 min. A second amplification was performed using Q5 polymerase (NEB) as follows: 72 °C 3 min, 98 °C 30 s, then 12 cycles of (98 °C 15 s, 60 °C 30 s, 72 °C 3 min), final extension at 72 °C 5 min. Libraries were purified using Vazyme DNA clean beads. The clean reads were mapped to the mouse genome (mm10) through STAR v2.2.0 with default settings. FeatureCounts software was used to collect reads located in the coding sequence (CDS) of each gene. CDS counts normalized to mappable reads were used for measuring gene expression. To compare gene expression between Zfp318-deficient and Zfp318-expressing MBCs across three different experimental settings, z-score of each gene was firstly calculated within each dataset to preclude batch and background bias, and then k-means clustering analysis was conducted using R software (v4.0.2). Gene set analysis was performed using GSEA software (v4.1.0). Expression profiles of all genes in the three settings were used as the input dataset and gene set c5.go.bp.v2023.1.Hs.symbols.gmt was used for enrichment analysis.

### QUANTIFICATION AND STATISTICAL ANALYSIS

For all relevant animal experiments, age- and sex-matched mice were randomly chosen to be included in different treatment groups. Each group typically had 3 to 5 animals, and 2 to 6 independent experiments were conducted for every assay. No blinding was necessary for animal experiments involved in this work. Except when noted otherwise, two-tailed Student’s *t* tests were used to compare endpoint means of different groups. Graphs were made in Prism (GraphPad).

---

**Figure S1.**
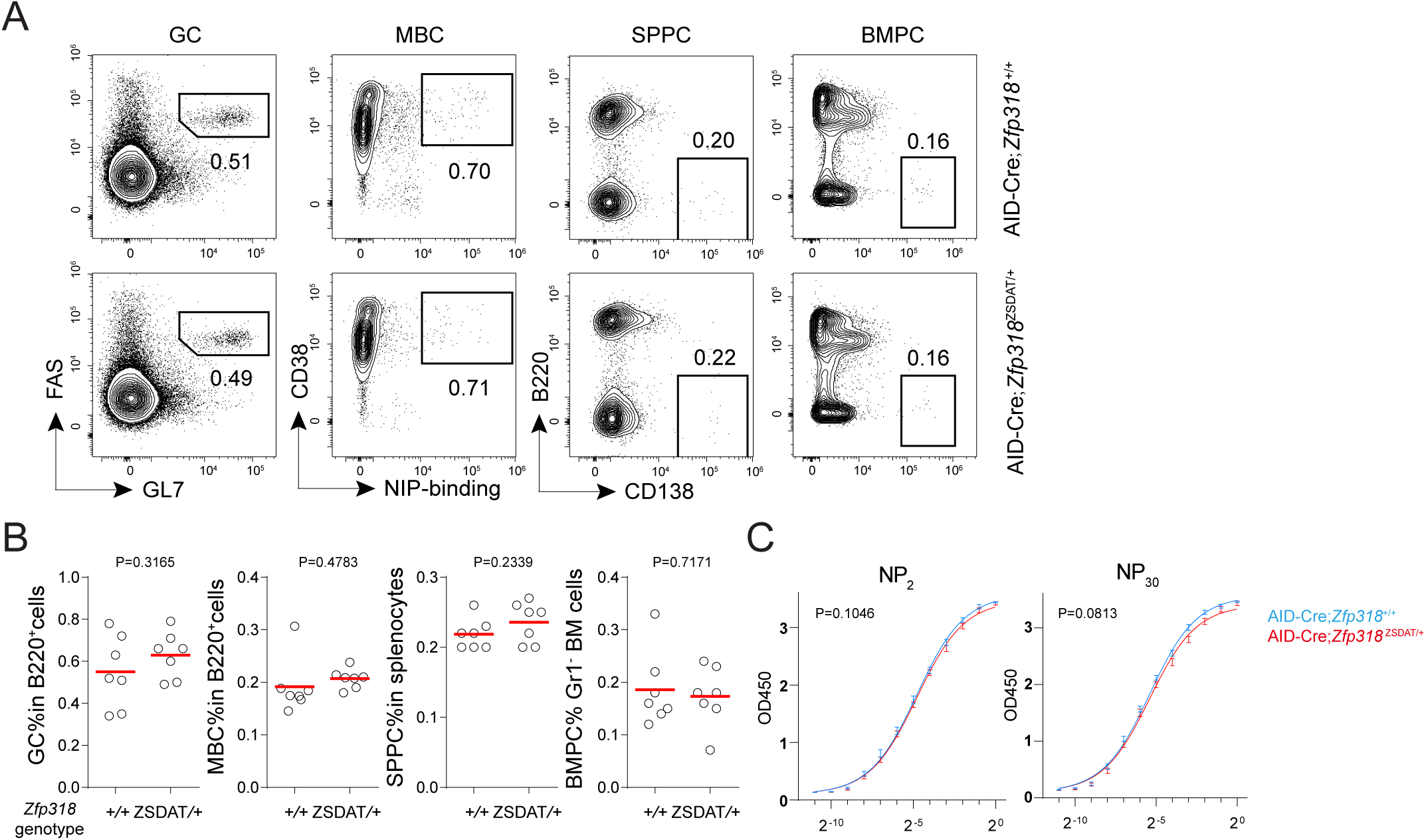
Loss of one functional *Zfp318* allele does not impair the primary response, related to Figure 1. **(A-C)** FACS profiles (**A**) and summary data (**B**) of GCs, SPPCs, BMPCs, serum NP_2_- and NP_30_-binding antibodies (**C**) in AID-Cre;*Zfp318*^+/+^ or AID-Cre;*Zfp318*^ZSDAT/+^mice on day 28 after NP-KLH immunization. In all scatter plots, each symbol is one mouse, and lines denote means. Data are pooled from 2 independent experiments. *P* values by *t* tests.

**Figure S2.**
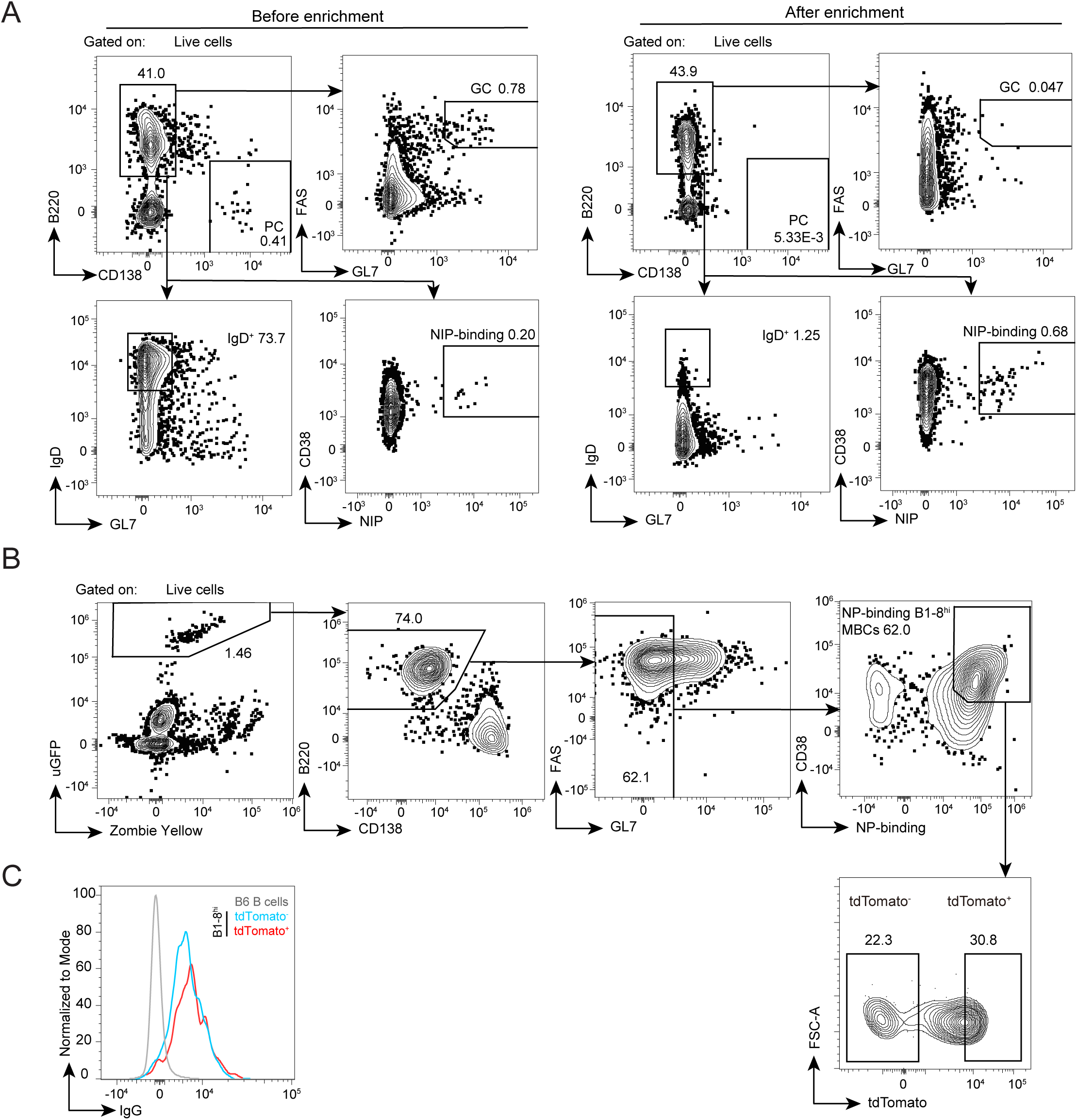
MBC enrichment for adoptive transfer assays, related to Figure 2. **(A)** FACS profiles of cells before (left) and after (right) MBC enrichment by magnetic depletion of CD4^+^ T cells, CD8^+^ T cells, IgD^+^ naïve B cells, GL7^+^ GC B cells, and CD138^+^ PCs followed by magnetic enrichment of CD19^+^ cells. **(B)** Gating strategy for sorting NP-binding B1-8^hi^ MBCs from donor mice of an uGFP-expressing AID-Cre;*Zfp318*^ZSDAT/+^ genotype. **(C)** Histogram overlay of IgG fluorescence of tdTomato^+^ and tdTomato^-^ B1-8^hi^ MBCs and B6 total B cells.

**Figure S3.**
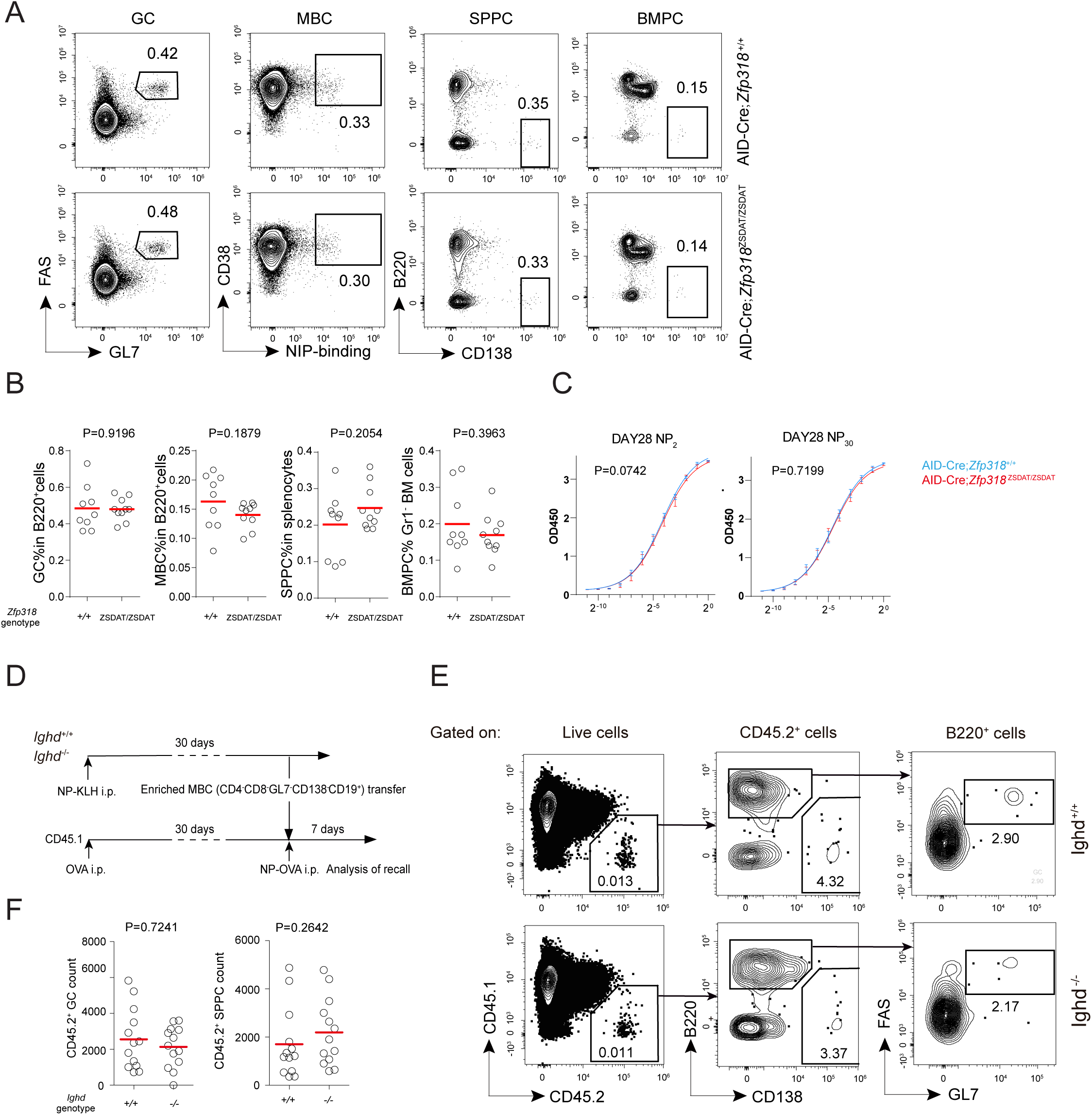
ZFP318 is not required for the primary response, and IgD is not required for MBC recall, related to Figure 3. **(A-C)** FACS profiles (**A**) and summary data (**B**) of GCs, SPPCs, BMPCs, serum anti-NP_2_ and anti-NP_30_ antibodies (**C**) in AID-Cre;*Zfp318*^+/+^ or AID-Cre;*Zfp318*^ZSDAT/ZSDAT^ mice on day 28 after NP-KLH immunization. In all scatter plots, each symbol is one mouse, and lines denote means. Data are pooled from 3 independent experiments. **(D-F)** Recall of MBCs from *Ighd*^+/+^ and *Ighd*^-/-^ mice. (**D)** Experimental design. **(E-F)** Representative FACS profiles (**E**) and summary data (**F**) of GCs and splenic PCs 7 days after re-challenge, as done according to (**D**). Each symbol indicating one mouse and lines denoting means. *P* values by *t* tests.

**Figure S4.**
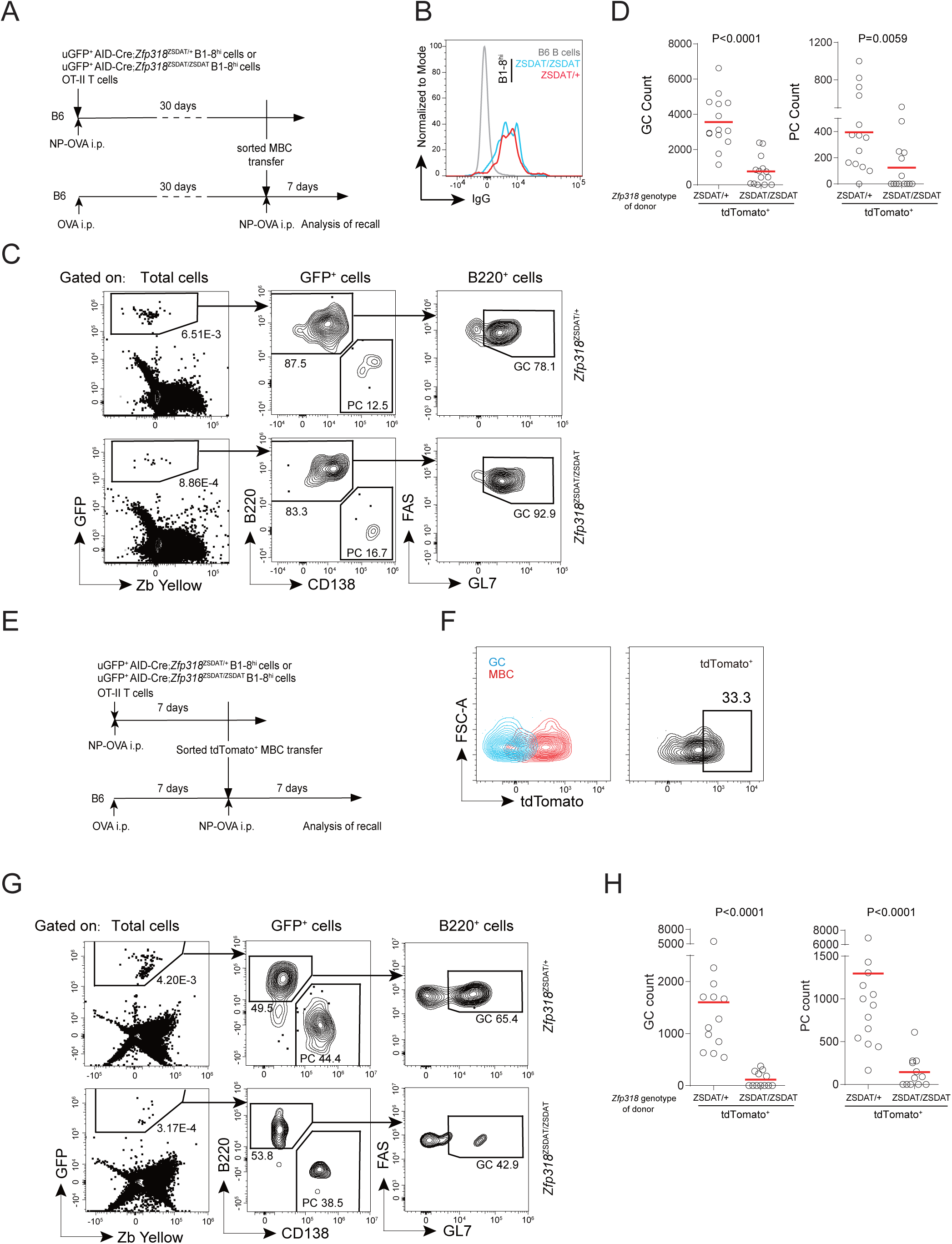
ZFP318 underlies MBC recallability, related to Figure 4. **(A-D)** Recall experiments utilizing adoptive transfer of GFP-expressing AID-Cre*;Zfp318*^ZSDAT/+^ or AID-Cre;*Zfp318*^ZSDAT/ZSDAT^ B1-8^hi^ cells isolated on day 30 after immunization. **(A)** Experimental design. Top 30% tdTomato-expressing MBCs were sort-purified for the transfer. **(B)** Histogram overlay of IgG fluorescence of tdTomato^+^ B1-8^hi^ MBCs from AID-Cre*;Zfp318*^ZSDAT/+^ or AID-Cre;*Zfp318*^ZSDAT/ZSDAT^ mice, with total B cells from B6 mice as control. **(C-D)** Gating strategy to identify donor-derived cells (**B**) and summary data (**C**) of GCs and splenic PCs derived from *Zfp318*^ZSDAT/+^ and *Zfp318*^ZSDAT/ZSDAT^ B1-8^hi^ MBCs 7 days after recall, as done according to (**A**). Data were pooled from 2 independent experiments, with each symbol indicating one mouse and lines denoting means. *P* values by Mann-Whitney *t* tests. **(E-H)** Recall experiments utilizing adoptive transfer of GFP-expressing AID-Cre*;Zfp318*^ZSDAT/+^ or AID-Cre;*Zfp318*^ZSDAT/ZSDAT^ B1-8^hi^ cells isolated on day 7 after immunization. **(E**) Experimental design. **(F**) The gating strategy to sort-purify tdTomato^+^ B1-8^hi^ MBCs. Left, an overlay of GCs and MBCs to show their difference in tdTomato fluorescence. Right, tdTomato^+^ gate for MBCs. **(G-H)** Gating strategy to identify donor-derived cells (**G**) and summary data (**H**) of GCs and splenic PCs derived from *Zfp318*^ZSDAT/+^ and *Zfp318*^ZSDAT/ZSDAT^ B1-8^hi^ MBCs 7 days after recall. Data were pooled from 2 independent experiments, with each symbol indicating one mouse and lines denoting means. *P* values by Mann-Whitney tests.

**Figure S5.**
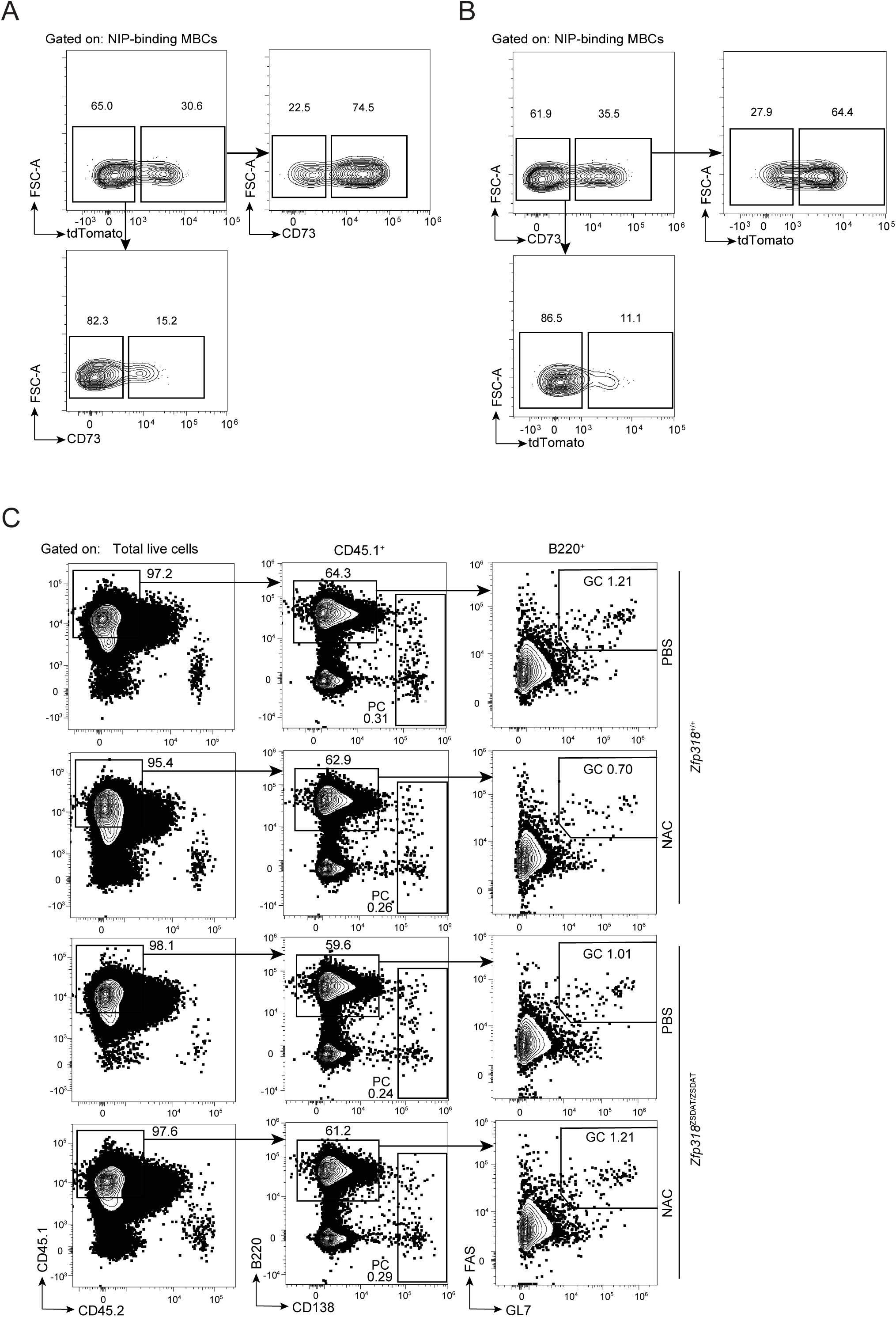
Control experiments related to Figure 2 and Figure 6. **(A-B)** Confirmation of CD73^+^ as a marker to enrich ZFP318-expressing MBCs for Fig. 2. **(A)** CD73 expression of tdTomato^+^ MBCs or tdTomato^-^ MBCs 28 days after NP-KLH immunization. **(B)** TdTomato expression of CD73^+^ MBCs or CD73^-^ MBCs 28 days after NP-KLH immunization. **(C)** Representative FACS profiles of CD45.1^+^ GCs and SPPCs, as in the experiment presented in Fig. 6.

**Figure S6.**
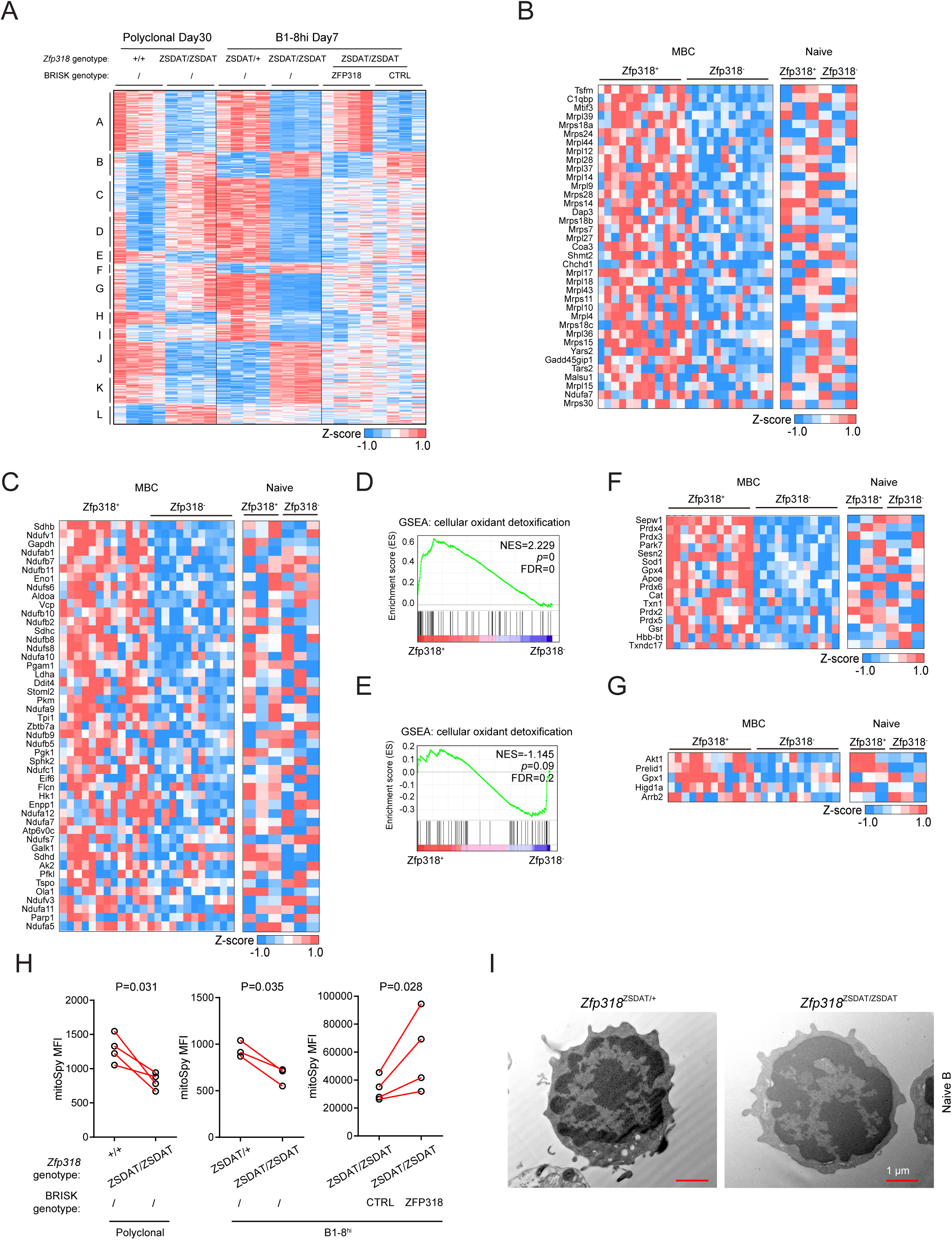
ZFP318 orchestrates a mitochondrial state conducive to recall, related to Figure 5. **(A-B)** Transcriptomic features of ZFP318-competent (3 conditions as described in the text: ZFP318-sufficient, ZFP318-heterozygous or ZFP318-induced, together noted as Zfp318^+^) MBCs compared to ZFP318-deficient (Zfp318^-^) MBCs. **(A)** K-means clustering analysis of differentially expressed genes (*P*<0.05 at least in one experimental setting) pooled from 3 experimental settings (see Fig. 5A-C) between Zfp318^+^ and Zfp318^-^ MBCs, identifying 12 clusters. **(B)** Heatmap of mitochondrial translation genes that are upregulated in Zfp318^+^ MBCs compared to Zfp318^-^ MBCs. Naïve B cells are used as controls. **(C)** Heatmap of genes involved in ATP synthesis that are upregulated in Zfp318^+^ MBCs compared to Zfp318^-^ MBCs. Naïve B cells are used as controls. **(D-E)** GSEA showing enrichment of cellular oxidant detoxification pathway in ZFP318^+^ compared to Zfp318^-^ MBCs (**D**) but not in naïve B cells (**E**). Normalized enrichment score (NES), *p* values, gene numbers and FDR are given in the graphs. **(F)** Heatmap showing genes involved in cellular oxidant detoxification that are upregulated in Zfp318^+^ MBCs compared to Zfp318^-^ MBCs. Naïve B cells are used as controls. **(G)** Heatmap showing genes involved in negative regulation of release of cytochrome c from mitochondria that are upregulated in Zfp318^+^ MBCs compared to Zfp318^-^ MBCs. Naïve B cells are used as controls. **(H)** Summary data of mitoSpy fluorescence of Zfp318^+^ and Zfp318^-^ MBCs in the 3 different experimental settings (Fig. 5A-C). Connected symbols are paired control and experimental groups from the same experiment. *P*-values by paired *t* tests. **(I)** Representative micrographs of sort-purified B1-8^hi^ naïve B cells of indicated types.

**Figure S7.**
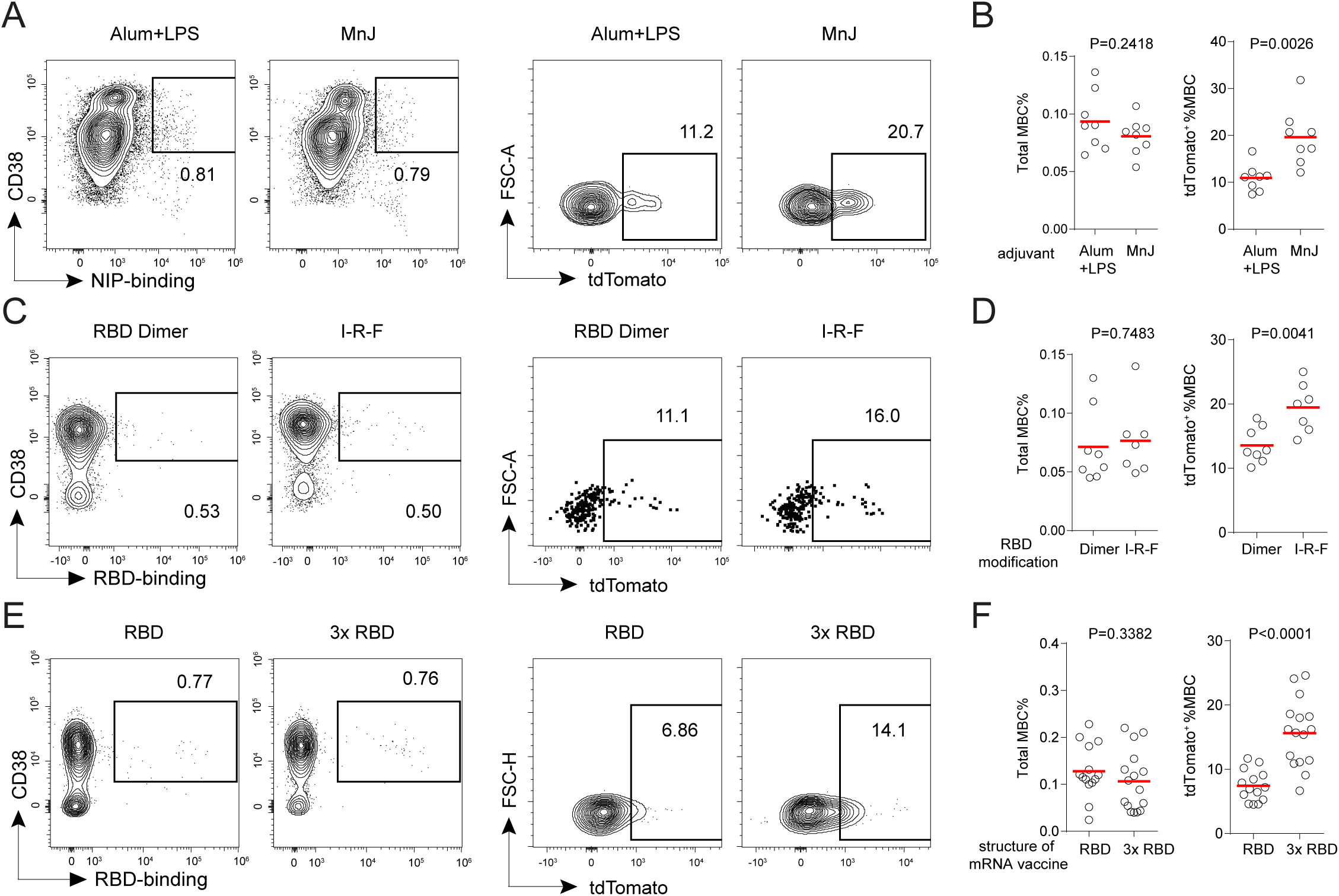
The abundance of ZFP318-expressing MBCs predicts the quality of prime-boost vaccination, related to Figure 7. **(A-B)** Comparison between two adjuvants. **(A)** Representative profiles of tdTomato fluorescence of MBCs 14 days after NP-KLH immunization adjuvanted with alum and LPS or MnJ, in *Zfp318*^ZSDAT/+^;AID-Cre mice. **(B)** Summary data of total MBCs as a fraction of splenic B cells or tdTomato^+^ MBCs as a fraction of total MBCs. Each symbol is one mouse, and lines denote means. Data are pooled from 4 independent experiments. **(C-D)** Comparison between two recombinant vaccine formula. **(C)** Representative profiles of tdTomato fluorescence of MBCs 14 days after immunization with a SARS-CoV2 RBD dimer or an IFN-RBD-Fc fusion, in *Zfp318*^ZSDAT/+^;AID-Cre mice. **(D)** Summary data of total MBCs as a fraction of splenic B cells or tdTomato^+^ MBCs as a fraction of total MBCs. Each symbol is one mouse, and lines denote means. Data are pooled from 2 independent experiments. **(E-F)** Comparison between two mRNA vaccine formula. **(E)** Representative profiles of tdTomato fluorescence of MBCs 21 days after immunization with mRNA vaccine expressing SARS-CoV2 RBD or SARS-CoV2 RBD trimer (3×RBD), in *Zfp318*^ZSDAT/+^;AID-Cre mice. **(F)** Summary data of total MBCs as a fraction of splenic B cells or tdTomato^+^ MBCs as a fraction of total MBCs. Each symbol is one mouse, and lines denote means. Data are pooled from 4 independent experiments. *P* values by *t* tests.

## Notes

### Summary of Updates

More data added to clarify ZFP318's function; Figures reorganized; Manuscript rewrote.

## References

1. Weisel, F., and Shlomchik, M. (2017). Memory B Cells of Mice and Humans. Annual Review of Immunology 35, 255–284. 10.1146/annurev-immunol-041015-055531.

2. Tarlinton, D., and Good-Jacobson, K. (2013). Diversity Among Memory B Cells: Origin, Consequences, and Utility. Science 341, 1205–1211. 10.1126/science.1241146.

3. Kurosaki, T., Kometani, K., and Ise, W. (2015). Memory B cells. Nature Reviews Immunology 15, 149–159. 10.1038/nri3802.

4. Dogan, I., Bertocci, B., Vilmont, V., Delbos, F., Mégret, J., Storck, S., Reynaud, C.-A., and Weill, J.-C. (2009). Multiple layers of B cell memory with different effector functions. Nat Immunol 10, 1292–1299. 10.1038/ni.1814.

5. Pape, K.A., Taylor, J.J., Maul, R.W., Gearhart, P.J., and Jenkins, M.K. (2011). Different B Cell Populations Mediate Early and Late Memory During an Endogenous Immune Response. Science 331, 1203–1207. 10.1126/science.1201730.

6. McHeyzer-Williams, L.J., Milpied, P.J., Okitsu, S.L., and McHeyzer-Williams, M.G. (2015). Class-switched memory B cells remodel BCRs within secondary germinal centers. Nat Immunol 16, 296–305. 10.1038/ni.3095.

7. Zuccarino-Catania, G.V., Sadanand, S., Weisel, F.J., Tomayko, M.M., Meng, H., Kleinstein, S.H., Good-Jacobson, K.L., and Shlomchik, M.J. (2014). CD80 and PD-L2 define functionally distinct memory B cell subsets that are independent of antibody isotype. Nature Immunology 15, 631–637. 10.1038/ni.2914.

8. Burton, D.R., Ahmed, R., Barouch, D.H., Butera, S.T., Crotty, S., Godzik, A., Kaufmann, D.E., McElrath, J.M., Nussenzweig, M.C., Pulendran, B., et al. (2012). A Blueprint for HIV Vaccine Discovery. Cell Host & Microbe 12. 10.1016/j.chom.2012.09.008.

9. Corti, D., and Lanzavecchia, A. (2013). Broadly Neutralizing Antiviral Antibodies. Immunology 31, 705–742. 10.1146/annurev-immunol-032712-095916.

10. Laidlaw, B.J., and Cyster, J.G. (2020). Transcriptional regulation of memory B cell differentiation. Nat Rev Immunol, 1–12. 10.1038/s41577-020-00446-2.

11. Inoue, T., and Kurosaki, T. (2023). Memory B cells. Nat. Rev. Immunol., 1–13. 10.1038/s41577-023-00897-3.

12. Shinnakasu, R., Inoue, T., Kometani, K., Moriyama, S., Adachi, Y., Nakayama, M., Takahashi, Y., Fukuyama, H., Okada, T., and Kurosaki, T. (2016). Regulated selection of germinal-center cells into the memory B cell compartment. Nature Immunology. 10.1038/ni.3460.

13. Wang, Y., Shi, J., Yan, J., Xiao, Z., Hou, X., Lu, P., Hou, S., Mao, T., Liu, W., Ma, Y., et al. (2017). Germinal-center development of memory B cells driven by IL-9 from follicular helper T cells. Nature immunology 18, 921–930. 10.1038/ni.3788.

14. Laidlaw, B.J., Schmidt, T.H., Green, J.A., Allen, C.D.C., Okada, T., and Cyster, J.G. (2017). The Eph-related tyrosine kinase ligand Ephrin-B1 marks germinal center and memory precursor B cells. J Exp Medicine 214, 639–649. 10.1084/jem.20161461.

15. Suan, D., Kräutler, N.J., Maag, J.L.V., Butt, D., Bourne, K., Hermes, J.R., Avery, D.T., Young, C., Statham, A., Elliott, M., et al. (2017). CCR6 Defines Memory B Cell Precursors in Mouse and Human Germinal Centers, Revealing Light-Zone Location and Predominant Low Antigen Affinity. Immunity 47, 1142–1153.e4. 10.1016/j.immuni.2017.11.022.

16. Ishizuka, M., Ohtsuka, E., Inoue, A., Odaka, M., Ohshima, H., Tamura, N., Yoshida, K., Sako, N., Baba, T., Kashiwabara, S., et al. (2016). Abnormal spermatogenesis and male infertility in testicular zinc finger protein Zfp318-knockout mice. Dev Growth Differ 58, 600–608. 10.1111/dgd.12301.

17. Enders, A., Short, A., Miosge, L.A., Bergmann, H., Sontani, Y., Bertram, E.M., Whittle, B., Balakishnan, B., Yoshida, K., Sjollema, G., et al. (2014). Zinc-finger protein ZFP318 is essential for expression of IgD, the alternatively spliced Igh product made by mature B lymphocytes. Proc National Acad Sci 111, 4513–4518. 10.1073/pnas.1402739111.

18. Pioli, P.D., Debnath, I., Weis, J.J., and Weis, J.H. (2014). Zfp318 Regulates IgD Expression by Abrogating Transcription Termination within the Ighm/Ighd Locus. J Immunol 193, 2546–2553. 10.4049/jimmunol.1401275.

19. Crouch, E.E., Li, Z., Takizawa, M., Fichtner-Feigl, S., Gourzi, P., Montaño, C., Feigenbaum, L., Wilson, P., Janz, S., Papavasiliou, F.N., et al. (2007). Regulation of AID expression in the immune response. J. Exp. Med. 204, 1145–1156. 10.1084/jem.20061952.

20. Zhang, Y., Meyer-Hermann, M., George, L.A., Figge, M.T., Khan, M., Goodall, M., Young, S.P., Reynolds, A., Falciani, F., Waisman, A., et al. (2013). Germinal center B cells govern their own fate via antibody feedback. J Exp Med 210, 457–464. 10.1084/jem.20120150.

21. Liu, D., Xu, H., Shih, C., Wan, Z., Ma, X., Ma, W., Luo, D., and Qi, H. (2015). T–B-cell entanglement and ICOSL-driven feed-forward regulation of germinal centre reaction. Nature 517, 214–218. 10.1038/nature13803.

22. Shlomchik, M.J. (2018). Do Memory B Cells Form Secondary Germinal Centers? Yes and No. Csh Perspect Biol 10, a029405. 10.1101/cshperspect.a029405.

23. McHeyzer-Williams, L.J., Dufaud, C., and McHeyzer-Williams, M.G. (2017). Do Memory B Cells Form Secondary Germinal Centers?: Impact of Antibody Class and Quality of Memory T-Cell Help at Recall. Csh Perspect Biol 10, a028878. 10.1101/cshperspect.a028878.

24. Pape, K.A., and Jenkins, M.K. (2017). Do Memory B Cells Form Secondary Germinal Centers?: It Depends. Csh Perspect Biol 10, a029116. 10.1101/cshperspect.a029116.

25. Roes, J., and Rajewsky, K. (1993). Immunoglobulin D (IgD)-deficient mice reveal an auxiliary receptor function for IgD in antigen-mediated recruitment of B cells. J Exp Medicine 177, 45–55. 10.1084/jem.177.1.45.

26. Taylor, J.J., Pape, K.A., and Jenkins, M.K. (2012). A germinal center– independent pathway generates unswitched memory B cells early in the primary response. J Exp Med 209, 597–606. 10.1084/jem.20111696.

27. Elgueta, R., Marks, E., Nowak, E., Menezes, S., Benson, M., Raman, V.S., Ortiz, C., O’Connell, S., Hess, H., Lord, G.M., et al. (2015). CCR6-Dependent Positioning of Memory B Cells Is Essential for Their Ability To Mount a Recall Response to Antigen. J Immunol 194, 505–513. 10.4049/jimmunol.1401553.

28. Glaros, V., Rauschmeier, R., Artemov, A.V., Reinhardt, A., Ols, S., Emmanouilidi, A., Gustafsson, C., You, Y., Mirabello, C., Björklund, Å.K., et al. (2021). Limited access to antigen drives generation of early B cell memory while restraining the plasmablast response. Immunity 54, 2005–2023.e10. 10.1016/j.immuni.2021.08.017.

29. Akkaya, M., Traba, J., Roesler, A.S., Miozzo, P., Akkaya, B., Theall, B.P., Sohn, H., Pena, M., Smelkinson, M., Kabat, J., et al. (2018). Second signals rescue B cells from activation-induced mitochondrial dysfunction and death. Nat Immunol 19, 871– 884. 10.1038/s41590-018-0156-5.

30. Purtha, W.E., Tedder, T.F., Johnson, S., Bhattacharya, D., and Diamond, M.S. (2011). Memory B cells, but not long-lived plasma cells, possess antigen specificities for viral escape mutants. J Exp Med 208, 2599–2606. 10.1084/jem.20110740.

31. Zhang, R., Wang, C., Guan, Y., Wei, X., Sha, M., Yi, M., Jing, M., Lv, M., Guo, W., Xu, J., et al. (2021). Manganese salts function as potent adjuvants. Cell Mol Immunol 18, 1222–1234. 10.1038/s41423-021-00669-w.

32. Wang, C., Guan, Y., Lv, M., Zhang, R., Guo, Z., Wei, X., Du, X., Yang, J., Li, T., Wan, Y., et al. (2018). Manganese Increases the Sensitivity of the cGAS-STING Pathway for Double-Stranded DNA and Is Required for the Host Defense against DNA Viruses. Immunity 48, 675–687.e7. 10.1016/j.immuni.2018.03.017.

33. Sun, S., Cai, Y., Song, T.-Z., Pu, Y., Cheng, L., Xu, H., Sun, J., Meng, C., Lin, Y., Huang, H., et al. (2021). Interferon-armed RBD dimer enhances the immunogenicity of RBD for sterilizing immunity against SARS-CoV-2. Cell Res, 1–13. 10.1038/s41422-021-00531-8.

34. Liang, Q., Wang, Y., Zhang, S., Sun, J., Sun, W., Li, J., Liu, Y., Li, M., Cheng, L., Jiang, Y., et al. (2022). RBD trimer mRNA vaccine elicits broad and protective immune responses against SARS-CoV-2 variants. Iscience 25, 104043. 10.1016/j.isci.2022.104043.

35. Dixon, S.J., and Olzmann, J.A. (2024). The cell biology of ferroptosis. Nat. Rev. Mol. Cell Biol., 1–19. 10.1038/s41580-024-00703-5.

36. Toyama, H., Okada, S., Hatano, M., Takahashi, Y., Takeda, N., Ichii, H., Takemori, T., Kuroda, Y., and Tokuhisa, T. (2002). Memory B Cells without Somatic Hypermutation Are Generated from Bcl6-Deficient B Cells. Immunity 17, 329–339. 10.1016/s1074-7613(02)00387-4.

37. Takemori, T., Kaji, T., Takahashi, Y., Shimoda, M., and Rajewsky, K. (2014). Generation of memory B cells inside and outside germinal centers. European Journal of Immunology 44, 1258–1264. 10.1002/eji.201343716.

38. Kaji, T., Ishige, A., Hikida, M., Taka, J., Hijikata, A., Kubo, M., Nagashima, T., Takahashi, Y., Kurosaki, T., Okada, M., et al. (2012). Distinct cellular pathways select germline-encoded and somatically mutated antibodies into immunological memory. J Exp Med 209, 2079–2097. 10.1084/jem.20120127.

39. Sokal, A., Chappert, P., Barba-Spaeth, G., Roeser, A., Fourati, S., Azzaoui, I., Vandenberghe, A., Fernandez, I., Meola, A., Bouvier-Alias, M., et al. (2021). Maturation and persistence of the anti-SARS-CoV-2 memory B cell response. Cell 184, 1201–1213.e14. 10.1016/j.cell.2021.01.050.

40. Burton, A.R., Guillaume, S.M., Foster, W.S., Wheatley, A.K., Hill, D.L., Carr, E.J., and Linterman, M.A. (2022). The memory B cell response to influenza vaccination is impaired in older persons. Cell Reports 41, 111613. 10.1016/j.celrep.2022.111613.

41. Chappert, P., Huetz, F., Espinasse, M.-A., Chatonnet, F., Pannetier, L., Silva, L.D., Goetz, C., Mégret, J., Sokal, A., Crickx, E., et al. (2022). Human anti-smallpox long-lived memory B cells are defined by dynamic interactions in the splenic niche and long-lasting germinal center imprinting. Immunity 55, 1872–1890.e9. 10.1016/j.immuni.2022.08.019.

42. Herzenberg, L., Black, S., Tokuhisa, T., and Herzenberg, L. (1980). Memory B cells at successive stages of differentiation. Affinity maturation and the role of IgD receptors. The Journal of experimental medicine 151, 1071–1087.

43. Mesin, L., Schiepers, A., Ersching, J., Barbulescu, A., Cavazzoni, C.B., Angelini, A., Okada, T., Kurosaki, T., and Victora, G.D. (2020). Restricted Clonality and Limited Germinal Center Reentry Characterize Memory B Cell Reactivation by Boosting. Cell 180, 92–106.e11. 10.1016/j.cell.2019.11.032.

44. Lau, A.W.Y., Turner, V.M., Bourne, K., Hermes, J.R., Chan, T.D., and Brink, R. (2020). BAFFR controls early memory B cell responses but is dispensable for germinal center function. J Exp Med 218, e20191167. 10.1084/jem.20191167.

45. Müller-Winkler, J., Mitter, R., Rappe, J.C.F., Vanes, L., Schweighoffer, E., Mohammadi, H., Wack, A., and Tybulewicz, V.L.J. (2020). Critical requirement for BCR, BAFF, and BAFFR in memory B cell survival. J Exp Med 218, e20191393. 10.1084/jem.20191393.

